# Ogg1 regulates context-dependent glial states in Alzheimer’s disease

**DOI:** 10.64898/2026.11.06.764857

**Authors:** Andreas Abentung, Teri Sakshaug, Paulina Schnur, Milena A. Egiazarian, Catarina Damião, Karoline Marie Aakre, Tobias Obermann, Mirta M. L. de Sousa, Nicolas Kunath, Magnar Bjørås, Katja Scheffler

**Affiliations:** Department of Clinical and Molecular Medicine Norwegian University of Science and Technology (NTNU), Trondheim, Norway; Department of Neuromedicine and Movement Science, Norwegian University of Science and Technology (NTNU), Trondheim, Norway; Central Administration, St Olavs Hospital, The University Hospital in Trondheim, Norway; Department of Neurology and Clinical Neurophysiology, St Olavs Hospital, The University Hospital in Trondheim, Norway; Department of Microbiology, Oslo University Hospital HF, Rikshospitalet and University of Oslo, Norway Rikshospitalet, Oslo University Hospital, Norway; Norwegian Health Association Dementia Research Center, Trondheim, Norway

## Abstract

Oxidative DNA damage is an early hallmark of Alzheimer’s disease (AD), but whether Ogg1, a key enzyme in oxidative DNA repair, contributes to disease pathogenesis remains unclear. Here, we show that Ogg1 exerts opposing effects under physiological and disease conditions. Ogg1 deficiency impaired memory and induced glial transcriptional alterations under physiological conditions, whereas in APP/PS1 mice it improved memory, reduced amyloid burden and attenuated AD-associated glial remodeling. These effects occurred without changes in global 8-oxo-dG levels and were accompanied by changes in chromatin accessibility and gene expression, supporting a non-canonical role for Ogg1 in transcriptional regulation. Single-nucleus transcriptomics revealed strong transcriptional responses to Ogg1 deficiency across all major glial cell types and showed that Ogg1-dependent glial transcriptional programs are strongly conserved in human AD. These findings identify Ogg1 as a context-dependent regulator of glial cell states and reveal an unexpected role in AD-associated glial remodeling.

## Introduction

Alzheimer’s disease (AD) is the most common cause of dementia, characterized by progressive cognitive decline and memory loss. Pathologically, AD is defined by the accumulation of amyloid-β (Aβ) plaques, hyperphosphorylated tau neurofibrillary tangles, synaptic dysfunction and neuronal loss^1^. Despite extensive efforts targeting amyloid-β and tau for AD treatment, clinical benefits have been limited, highlighting the need for novel therapeutic targets that affect upstream regulatory mechanisms rather than downstream pathological symptoms^2^.

Aging is the strongest risk factor for AD and is accompanied by a progressive accumulation of DNA damage, leading to somatic mutations, transcriptional dysregulation, and cognitive decline in the human brain^3,4^. Consistent with the brain’s vulnerability to genomic instability, increased DNA damage and impaired DNA repair have been implicated in premature aging and neurological phenotypes in both humans and rodents^5,6^. Among the different DNA damage types, oxidative lesions are particularly prominent in the brain, and neurons and glial cells are highly susceptible to DNA damage due to high metabolic demand, chronic oxidative stress, and limited regenerative capacity^7,8^. Base excision repair (BER) is the primary pathway responsible for resolving oxidative lesions, with 8-oxoguanine DNA glycosylase-1 (Ogg1) serving as the key enzyme initiating repair of the pre-mutagenic lesion 8-oxoguanine^9^.

Previous studies have suggested that Ogg1-mediated DNA repair plays an important role in AD pathogenesis. A study using the AD APP NL-G-F knock-in mouse model has shown that Ogg1 deficiency accelerates age-dependent oxidative DNA damage and microgliosis^10^. In humans, an *OGG1* polymorphism is associated with AD compromising enzymatic activity and cellular resistance to oxidative stress^11^.

Beyond its canonical role in maintaining genome integrity, Ogg1 has emerged as a key regulator of gene transcription through epigenetic mechanisms that appear largely independent of DNA repair^12,13^. Ogg1 was shown to interact with histone deacetylase 1 (HDAC1), a central modulator of histone acetylation and chromatin accessibility, whose brain-specific loss leads to altered gene expression and age-dependent cognitive decline^14^. Moreover, DNA glycosylases, including Ogg1, have been implicated in shaping DNA methylation and Polycomb-repressive complex 2 (PRC2)-associated histone modifications at genes involved in higher human brain function^15^. Together, these findings implicate Ogg1 in epigenetic regulation in the brain, with the potential to influence gene-regulatory programs in aging and disease. However, the extent to which Ogg1-dependent epigenetic regulation contributes to transcriptional changes and pathogenesis in AD remains unknown.

Here, we combined bulk transcriptomic profiling, chromatin accessibility analysis, and single-nucleus RNA sequencing (snRNA-seq) in wildtype and Ogg1-deficient mice as well as APP/PS1 mice with and without Ogg1 deficiency. We find that loss of Ogg1 alone impairs long-term memory to a similar extent as observed in APP/PS1 mice. Notably, Ogg1 deficiency in the context of AD prevents this memory impairment, coinciding with a lower amyloid burden. Bulk RNA sequencing and chromatin accessibility profiling point to dysregulated inflammatory signaling as a major driver of this phenotype. SnRNA-seq further reveals that Ogg1 loss prevents AD-associated transcriptional dysregulation in immune response, synaptic signaling, and myelination across microglia, astrocytes, and oligodendrocytes. Importantly, these transcriptional changes largely recapitulate the gene expression alterations observed in human AD patients and are accompanied by widespread changes in chromatin accessibility across glial cell types. Together, our findings reveal a previously unrecognized role of Ogg1 in mediating cell type-specific glial gene regulatory programs in AD.

## Results

### Ogg1 deficiency prevents cognitive decline and inflammatory dysregulation in APP/PS1 mice

To determine whether OGG1 is implicated in human AD, we analyzed *OGG1* expression in post-mortem medial temporal lobe tissue from patients with AD. *OGG1* expression was increased at both the mRNA and protein level in AD patients relative to healthy donors (Supp.Fig1a, i and ii). These findings motivated us to investigate whether Ogg1 functionally contributes to AD pathogenesis. We therefore generated APP/PS1×*Ogg1^-/-^* (AO) mice by crossing APP/PS1 mice with *Ogg1* knockout mice^16,17^. All experiments were performed at 6 months of age. To assess the impact of Ogg1 deficiency on amyloid pathology, we quantified amyloid-β (Aβ) plaque deposition in the cortex and hippocampus of APP/PS1 and AO mice (Fig.1a). No significant differences were observed in plaque number, plaque-covered area, or plaque size in either brain region (Fig.1b; Supp.Fig1b, i and ii). Furthermore, we examined the levels of the major amyloid-β species, Aβ42 and Aβ40, in whole mouse-brain lysates by ELISA. AO mice showed significantly reduced levels of insoluble Aβ42 and Aβ40 (Fig.1 c) demonstrating that Ogg1 deficiency attenuates amyloid pathology. In contrast, levels of soluble Aβ42 and Aβ40 did not differ between APP/PS1 and AO mice (Supp.Fig1c). To determine whether reduced insoluble Aβ accumulation translates into functional benefits, we subjected mice to the open field test to evaluate anxiety-like behavior and to the novel object location (NOL) paradigm to assess short-term (1 h) and long-term (24 h) spatial memory. No major differences were observed across genotypes in total distance traveled, mean speed, time spent in the center, or number of center entries (Supp.Fig2a and b), indicating the absence of an anxiety-like phenotype. In the 1h memory retention test, all genotypes exhibited similar exploration of the novel object, indicating intact short-term spatial memory (Supp.Fig2c). In contrast, during the 24 h memory retention test, both *Ogg1*^-/-^ and APP/PS1 mice displayed impaired spatial memory, spending significantly less time exploring the novel object location than WT mice (Fig.1d). Thus, Ogg1 deficiency alone was sufficient to impair long-term spatial memory under physiological conditions. Notably, Ogg1 deficiency rescued the memory deficit observed in APP/PS1 mice, as AO mice performed comparably to WT animals in exploring the novel object location. Consistent with this, *Ogg1^-/-^* and APP/PS1 mice showed no preference for the novel location based on head entries whereas AO mice displayed a significant preference for the novel location (Fig.1d). Object-specific exploration and head entry numbers during training (Supp.Fig2d and e), as well as total exploration time during the 24 h test (Supp.Fig2f), were comparable across all genotypes.

**Figure 1.**
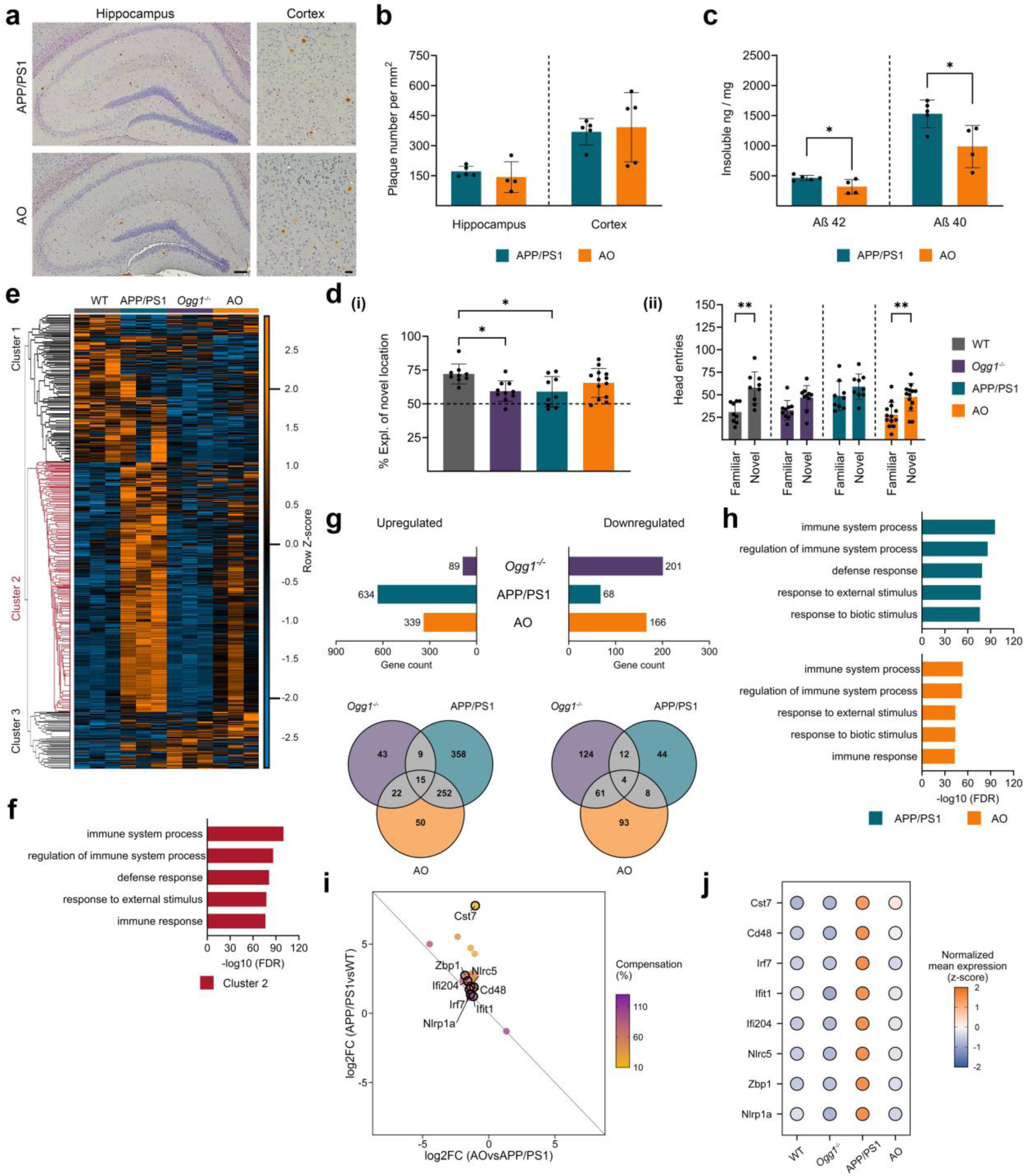
Ogg1 deficiency reduces amyloid burden, preserves long-term memory, and attenuates inflammatory transcriptional changes in APP/PS1 mice. (a) Representative images of immunohistochemically stained amyloid-β plaques in the hippocampus and cortex of APP/PS1 and AO mice. Scale bars: 100 µm (hippocampus) and 20 µm (cortex). (b) Quantification of plaque number in the brain of APP/PS1 and AO mice. (c) Analysis of insoluble Aβ42 and Aβ40 in whole-brain extracts using ELISA (APP/PS1, n = 5; AO, n = 4; for both measurements). Data are presented as mean ± SEM, n values refer to the number of mice, *p < 0.05. (d) (i) Long-term spatial memory was assessed using the novel object location (NOL) paradigm 24 h after training (WT, n = 9; *Ogg1^-/-^*, n = 10; APP/PS1, n = 9; AO, n = 13). (ii) Head entries directed toward the familiar and novel object during the 24 h NOL test in WT, *Ogg1^-/-^*, APP/PS1, and AO mice. Data are presented as mean ± SEM, n values refer to the number of mice, *p < 0.05. (e) Heatmap over all statistically significantly differentially expressed genes (DEGs) (|log2 fold change| ≥ 0.5, p < 0.05) in the hippocampus of mice, clustered using Euclidean distance and average linkage. Values are FPKMs, z-scored by row revealing three different gene clusters. (f) Gene ontology (GO) analysis of cluster 2 genes. Bar charts show the top five biological process terms. (g) Bar graphs illustrate the number of DEGs (|log₂FC| ≥ 0.5, p < 0.05) for *Ogg1^-/-^*, APP/PS1 and AO when compared to WT. Venn diagrams indicate the unique downregulated and upregulated genes for each genotype. (h) Enriched GO biological process terms for unique upregulated genes in either APP/PS1 or AO. (i) Scatter plot showing the relationship between gene expression changes in APP/PS1 versus WT and AO versus APP/PS1 mice. Each dot represents a preventative differentially expressed gene (prevDEG). The color gradient indicates the compensation rate (%), calculated as the ratio of the log₂ fold changes in AO versus APP/PS1 to that in APP/PS1 versus WT, with higher values indicating greater prevention of AD-associated transcriptional dysregulation. Black outlines indicate selected prevDEGs highlighted in the manuscript. (j) Dot plot showing z-scored mean expression (FPKM, scaled per gene across genotypes) of selected bulk RNA-seq prevDEGs in all genotypes.

Given the canonical role of Ogg1 in repairing 8-oxo-dG lesions, we next asked whether these phenotypic effects were associated with altered oxidative DNA damage. Liquid chromatography-tandem mass spectrometry (LC-MS/MS) measurements of hippocampal and cortical 8-oxo-dG levels revealed no differences between WT, *Ogg1^-/-^*, APP/PS1, and AO (Supp.Fig3, i and ii), indicating that global oxidative DNA damage does not accumulate across genotypes at this stage of disease.

These findings suggest that the impact of Ogg1 on AD pathology may extend beyond its established DNA repair function. To explore potential transcriptional mechanisms, we performed bulk RNA sequencing (bRNA-seq) of hippocampal tissue from all genotypes. Principal component analysis (PCA) showed consistent clustering of biological replicates but clear genotype separation with AO being more similar to WT than APP/PS1 (Supp.Fig4a). Hierarchical clustering of differentially expressed genes (DEGs) identified three distinct gene clusters, with cluster 2 exhibiting a unique expression profile in APP/PS1 that was largely absent in AO (Fig.1e). Gene ontology (GO) analysis of this cluster revealed a strong enrichment for immune response-related pathways (Fig.1f), whereas cluster 1 was associated with development-related pathways and no significant enrichment was detected for cluster 3 (Supp.Fig4b).

We next compared unique and shared DEGs across genotypes relative to WT (Fig.1g). APP/PS1 exhibited predominantly upregulated genes, whereas *Ogg1^-/-^* showed largely downregulated genes. Interestingly, the transcriptional profile of AO differed from that of *Ogg1^-/-^*, indicating that the effects of Ogg1 loss on gene expression are influenced by the presence of amyloid pathology. GO analysis of upregulated genes in APP/PS1 and AO mice identified similar immune-related pathways, although enrichment was reduced in AO mice, suggesting that Ogg1 deficiency attenuates inflammatory gene expression in the context of AD (Fig.1h). GO analysis of downregulated genes in *Ogg1^-/-^* and AO revealed enrichment of developmental processes (Supp.Fig4c).

To identify genes whose AD-associated dysregulation was prevented by Ogg1 deficiency, we intersected DEGs from the APP/PS1 vs WT and AO vs AD comparisons. Genes that changed significantly in opposite directions were defined as preventative DEGs (prevDEGs; Fig.1i). We quantified the extent of transcriptional compensation for each prevDEG by calculating a compensation rate (%), representing the proportion of the AD-associated expression change prevented by Ogg1 deficiency. This analysis identified 31 prevDEGs, including genes implicated in AD pathogenesis through their roles in microglial activation^18^, interferon signaling^19^, and antigen presentation^20^ (Fig.1j). Collectively, these findings suggest that Ogg1 deficiency partially reverses AD-associated inflammatory transcriptional programs.

### Ogg1 deficiency prevents AD-associated chromatin accessibility changes in APP/PS1 mice

To investigate the epigenetic mechanisms underlying the prevention of AD-associated transcriptional changes in AO mice, we performed Assay for Transposase-Accessible Chromatin using sequencing (ATAC-seq) on hippocampal tissue across all genotypes. PCA of the ATAC-seq count matrix showed tight clustering of biological replicates across all genotypes, confirming high data quality and reproducibility (Supp.Fig5a). Analysis of chromatin accessibility around transcription start sites (TSS) revealed largely comparable ATAC-seq read intensities across genotypes, indicating no major global changes in chromatin accessibility surrounding TSS (Supp.Fig5b).

We next examined genotype-specific and shared differentially accessible regions (DARs) relative to WT (Fig.2a). To ensure robustness, only DARs detected in at least two out of three biological replicates were considered for downstream analyses. APP/PS1 mice exhibited the largest number of DARs, with the majority showing increased accessibility and a substantial overlap with DARs identified in AO. In contrast, *Ogg1^-/-^* exhibit mainly DARs with reduced accessibility, 42% of which were also observed in AO. Most accessible chromatin regions were located within intronic and distal intergenic regions, followed by promoter regions (Supp.Fig5c). Genotype-specific DARs revealed that *Ogg1^-/-^* showed the lowest percentage of promoter-associated DARs, whereas APP/PS1 and AO showed broadly similar genomic distribution (Fig.2b).

Motif enrichment analysis of genotype-specific chromatin accessibility changes, integrating gained and lost DARs, revealed a prominent enrichment of immune-associated transcription factor motifs in APP/PS1 mice. Among the top enriched motifs was PU.1, a key regulator of microglial identity and function in AD^21^ (Fig.2d). In contrast, Ogg1-specific DARs showed enrichment for only a limited set of motifs, including ZAC1 and NFAT, transcription factors implicated in neuronal progenitor differentiation^22^ and activity-dependent gene expression^23^, respectively (Supp.Fig5d). AO-specific DARs were enriched for motifs associated with transcription factors involved in neuronal differentiation^24^, metabolic regulation, and synaptic plasticity^25^ (Supp.Fig5e).

Next, we asked whether Ogg1 deficiency counteracts AD-associated changes in chromatin accessibility. Using the same approach as for prevDEGs, we identified preventative differentially accessible regions (prevDARs), defined as regions exhibiting significant accessibility changes in opposite directions in the APP/PS1 versus WT and AO versus APP/PS1 comparisons (Fig.2 e). We identified 83 prevDARs, indicating that loss of Ogg1 partially prevents AD-associated changes in chromatin accessibility. Notably, several prevDARs mapped to the Firre locus, which encodes a long non-coding RNA involved in higher-order chromatin organization^26^, highlighting it as a recurrent locus of preventative chromatin accessibility changes (Supp.Fig5f). Consistent with this observation, chromatin accessibility across the Firre gene body was increased in APP/PS1 mice but remained at WT-like levels in AO mice (Fig.2 f). These findings raise the possibility that Ogg1-dependent regulation of Firre contributes to the broader chromatin remodeling associated with AD pathology.

**Figure 2.**
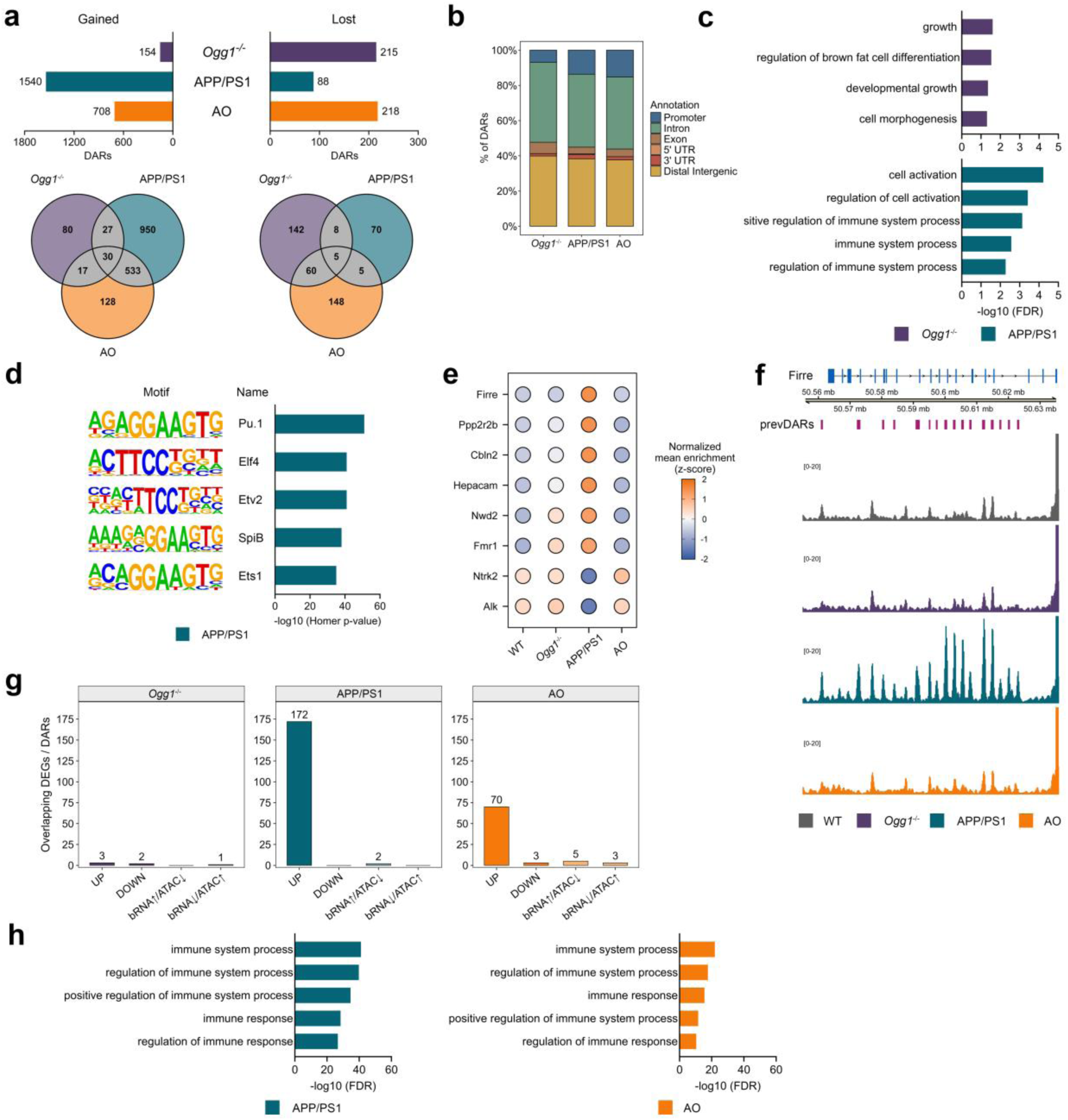
Ogg1 deficiency prevents immune-associated chromatin accessibility changes in APP/PS1 mice. (a) Bar graphs showing the number of differentially accessible regions (DARs) identified in *Ogg1^-/-^*, APP/PS1, and AO mice relative to WT (|log_2_ fold change| ≥ 1, p < 0.05). Venn diagrams depict unique gained and lost DARs for each genotype. (b) Genomic annotation of genotype-specific unique DARs. Prom, promoter; Dist, distal intergenic; Int, intron; Ex, exon; UTR, untranslated region. (c) Enriched gene ontology (GO) biological process terms associated with genotype-specific unique DARs in *Ogg1^-/-^*and APP/PS1 mice. Gained and lost DARs were analyzed together. (d) Transcription factor (TF) motif enrichment analysis of APP/PS1-specific unique DARs. Gained and lost DARs were analyzed together. (e) Dot plot showing z-scored mean accessibility of prevDAR-associated genes across genotypes (peaks averaged per gene). (f) Genome browser view of the Firre locus showing genotype-dependent differences in chromatin accessibility. PrevDARs are indicated above the tracks. (g) Bar graphs showing the overlap between differentially expressed genes (bulk RNA-seq) and genes associated with differentially accessible regions across genotypes, stratified by direction of regulation. (h) GO biological process enrichment analysis of genes showing concordant increases in chromatin accessibility and gene expression.

Because bRNA-seq and ATAC-seq independently identified immune-related pathways as the most prominently affected biological processes, we next asked whether transcriptional and chromatin alterations were coordinated. Integration of both datasets identified 172 genes with concordant increases in gene expression and chromatin accessibility in APP/PS1 mice, compared to only 70 genes in AO mice (Fig.2g). These genes were strongly enriched for immune-related processes in APP/PS1 mice, whereas this enrichment was reduced in AO mice (Fig.2h). This suggests that coordinated transcriptional and epigenetic remodeling underlies AD-associated inflammatory activation.

### SnRNA-seq reveals context-dependent neuronal transcriptional responses to Ogg1 loss

Given that AD-associated gene expression changes are highly cell type-specific^27,28^, we performed snRNA-seq of hippocampal tissue across all genotypes. Quality control metrics were comparable across biological replicates and genotypes and indicated robust data quality (Supp.Fig6). Uniform Manifold Approximation and Projection (UMAP) analysis resolved 47,715 nuclei into distinct clusters corresponding to the major hippocampal cell populations, as defined by established marker genes (Fig.3a and b). Cell-type proportions were comparable across genotypes and replicates, indicating no differences in overall cellular composition (Supp.Fig7a and b).

**Figure 3.**
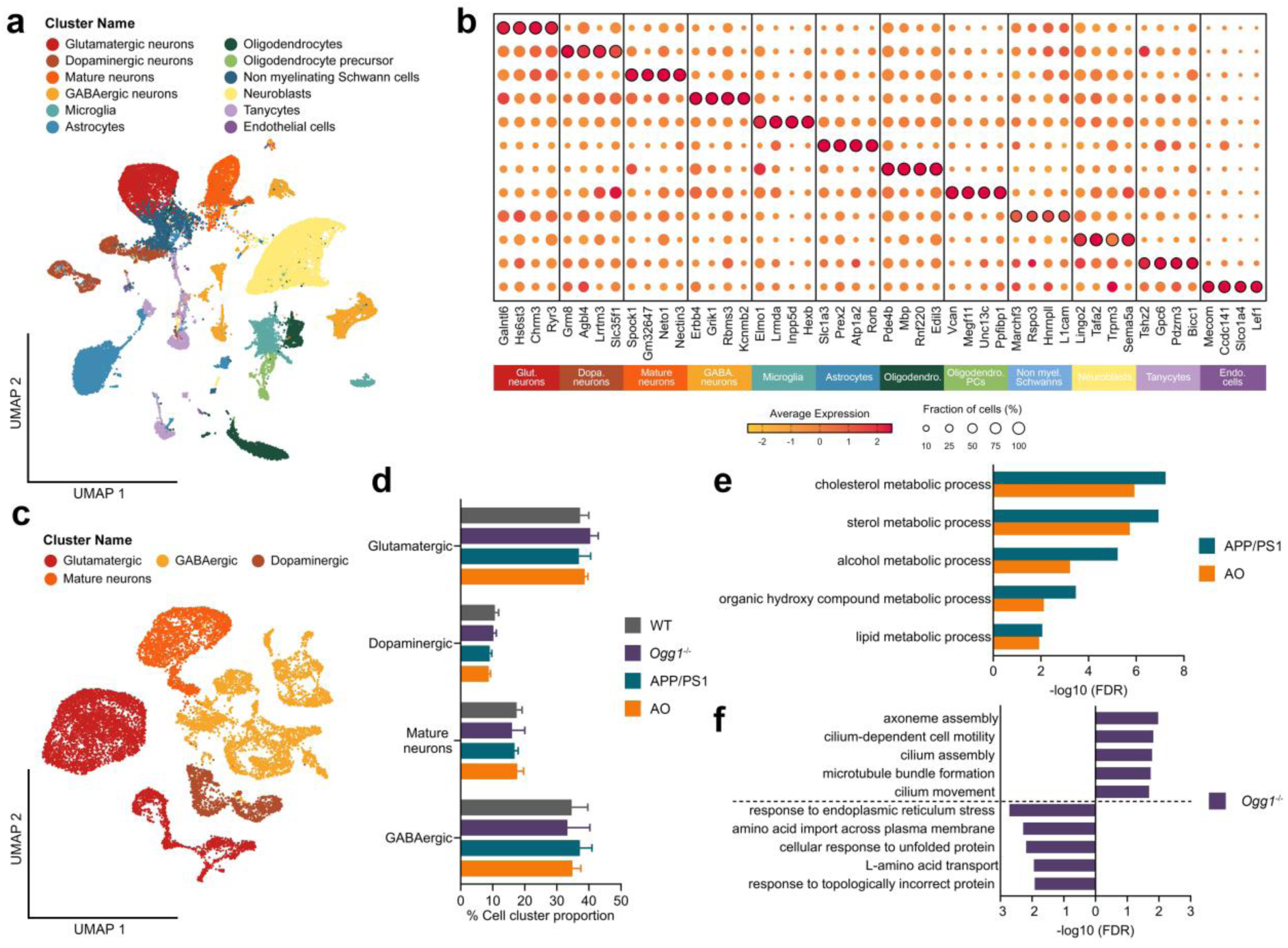
Ogg1 deficiency shapes neuronal gene expression in a context-dependent manner. (a) UMAP projection of single-nucleus RNA sequencing (snRNA-seq) data showing all identified cell clusters across 47,715 individual nuclei. Clusters were defined based on transcriptomic similarity and annotated according to established cell-type-specific marker genes. (b) Dot plot of four representative marker genes per cell cluster used for cell-type annotation. Dot size indicates the proportion of nuclei expressing each gene, while the color gradient reflects the average expression level. Outlined dots indicate the corresponding marker genes for each cell cluster. (c) UMAP visualization of snRNA-seq data showing all neuronal cell clusters. (d) Percentage distribution of nuclei among the four neuronal clusters: glutamatergic, GABAergic, dopaminergic, and mature neurons across all genotypes. (e) Enriched gene ontology (GO) biological process terms shared between APP/PS1 and AO for genotype-specific upregulated differentially expressed genes (DEGs) in glutamatergic neurons. ^(f)^ Enriched GO biological process terms for unique upregulated and downregulated DEGs in *Ogg1^-/-^* GABAergic neurons.

To define the cell type-specific effects of Ogg1 deficiency, we first focused on neuronal populations identified by snRNA-seq. UMAP analysis resolved distinct neuronal subclusters, including glutamatergic, GABAergic, dopaminergic, and mature neurons (Fig.3c). The relative abundance of these neuronal clusters was comparable across genotypes, indicating preserved neuronal subtype composition (Fig.3d). Differential expression analysis identified similar numbers of up- and downregulated genes across genotypes, however, changes were largely genotype-specific, with most DEGs unique to each genotype (Supp.Fig7, c - f). Despite limited overlap at the gene level, GO enrichment analysis identified shared pathways related to cholesterol and sterol metabolism among upregulated genes in glutamatergic neurons from APP/PS1 and AO mice (Fig.3e). Notably, these processes are tightly linked to AD pathology^29^ and their enrichment was attenuated in AO. In *Ogg1^-/-^*mice, upregulated genes in GABAergic neurons were enriched for axoneme and cilium assembly pathways, whereas downregulated genes were associated with responses to endoplasmic reticulum stress (Fig.3f). Together, these findings indicate that the effects of Ogg1 deficiency on neuronal gene expression are context-dependent and differ between physiological conditions and AD.

### Ogg1 deficiency suppresses AD-associated microglial activation

The attenuation of AD-associated inflammatory pathways in AO mice implicated microglia as key mediators of Ogg1-dependent effects in AD^30^. Consistent with this, snRNA-seq revealed extensive transcriptional remodeling across microglial populations (Fig.4a). UMAP analysis resolved two clusters corresponding to activated microglia (Act.1 and Act.2), alongside multiple homeostatic and intermediate microglial populations (Mic.3-Mic.12), which were functionally annotated using GO enrichment analysis. Notably, Act.1 was predominantly observed in APP/PS1 mice but largely prevented in AO mice, suggesting that Ogg1 deficiency suppresses the development of this activated microglial state in AD (Fig.4b). Interestingly, Act.2 was present across all genotypes and significantly more abundant in *Ogg1^-/-^*, indicating that distinct activated microglial programs arise following Ogg1 loss under both physiological conditions and in AD.

**Figure 4.**
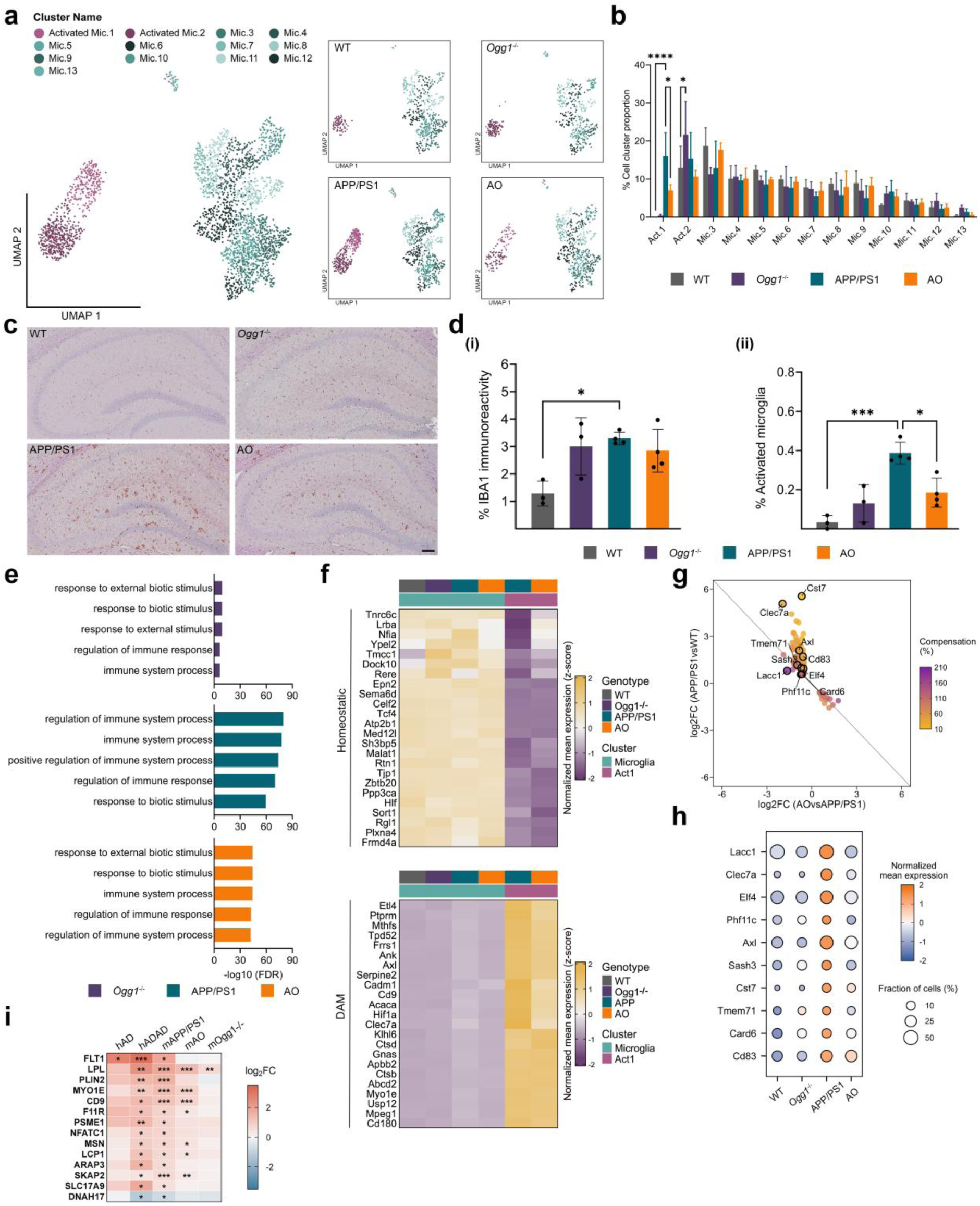
Ogg1 deficiency suppresses disease-associated microglial activation and inflammatory gene expression in APP/PS1 mice. (a) UMAP representation of combined- and genotype specific microglial nuclei derived from snRNA-seq, revealing subclusters defined by transcriptomic similarity and functional annotation based on gene ontology (GO) biological process enrichment. (b) Percentage distribution of nuclei among the microglial subclusters across all genotypes. Data are presented as mean ± SEM, n values refer to the number of mice, *p < 0.05, ****p < 0.0001. (c) Representative images of IBA1 immunostaining in the hippocampus across all genotypes. Scale bar: 100 µm. (d) Quantification of (i) hippocampal IBA1 immunoreactivity and (ii) proportion of activated hippocampal IBA1 positive microglia in WT, *Ogg1^-/-^*, APP/PS1 and AO mice. Data are presented as mean ± SEM, n values refer to the number of mice, *p < 0.05, **p < 0.01, ***p < 0.001. (e) GO biological process enrichment of genotype-specific upregulated genes in the microglial cluster of all genotypes. (f) Heatmap showing the normalized mean expression (z-score) of homeostatic and disease-associated microglia (DAM) genes, as reported by Keren-Shaul et al.^35^, across genotypes and microglial subclusters in our dataset. (g) Scatter plot showing the relationship between gene expression changes in APP/PS1 versus WT and AO versus APP/PS1 microglia. Each dot represents a preventative differentially expressed gene (prevDEG). The color gradient indicates the compensation rate (%), calculated as the ratio of the log₂ fold changes in AO versus APP/PS1 to that in APP/PS1 versus WT, with higher values indicating greater prevention of AD-associated transcriptional dysregulation. Genes outlined in black indicate selected prevDEGs. (h) Dot plot depicting normalized mean expression of prevDEGs across genotypes in microglia. Dot size reflects the percentage of nuclei expressing the indicated gene. (i) Comparison of significantly DEGs (|log_2_ fold change| ≥ 0.5, p < 0.05) in microglia from our mouse snRNA-seq dataset and human parietal cortex microglia from sporadic AD (hAD) and autosomal dominant AD (hADAD; APP and PSEN1) reported by Brase et al.^33^. Shown are genes significantly dysregulated in hADAD (Benjamini–Hochberg-adjusted p < 0.05) with the same direction of change in APP/PS1 mice. Color indicates log₂ fold changes relative to WT mice (mouse) or neuropathology-free controls (human). Genes highlighted in bold represent prevDEGs. Asterisks indicate genes that are significantly differentially expressed (*p < 0.05, **p < 0.01, ***p < 0.001; Benjamini–Hochberg-adjusted p for human, nominal p for mouse comparisons).

To determine whether the microglial states were reflected at the tissue level, we performed immunostaining for IBA1 (ionized calcium-binding adapter molecule 1). In the hippocampus, total IBA1 immunoreactivity was significantly increased in APP/PS1 mice relative to WT mice and showed a similar, albeit non-significant, trend in *Ogg1^-/-^* and AO mice (Fig.4c and d i). In the cortex, IBA1 immunoreactivity was significantly elevated in APP/PS1, *Ogg1^-/-^*, and AO mice (Supp.Fig8 a and b i). Because activated microglia exhibit enlarged somata and a less ramified morphology, we next quantified microglial activation by calculating the cell body-to-cell size ratio of IBA1-positive cells^31^. Activated microglia were more abundant in APP/PS1 mice but significantly reduced in AO mice in both the hippocampus and cortex (Fig.4d ii and Supp.Fig8 b ii). These findings closely mirrored the relative abundance of the Act.1 microglial subcluster identified by snRNA-seq. In addition, *Ogg1^-/-^* mice displayed a modest increase in activated microglia relative to WT mice, consistent with the expansion of the Act.2 microglial population.

Microglia-specific DEGs revealed a predominance of upregulated genes across all genotypes, with APP/PS1 mice exhibiting the largest number, followed by AO and *Ogg1^-/-^* mice (Supp.Fig8c). While APP/PS1 and AO shared a substantial proportion of DEGs, each genotype also exhibited a distinct transcriptional profile. Genotype-specific upregulated genes in APP/PS1 mice were enriched for immune- and inflammation-related biological processes, an effect markedly reduced in AO, but already evident in *Ogg1^-/-^* mice (Fig.4e). These findings indicate that Ogg1 deficiency promotes microglial inflammatory programs in the absence of pathology but suppresses them in the AD context.

To further define the microglial subclusters, we examined the expression of previously published disease-associated microglia (DAM) gene signatures^32,33^. Whereas the combined Mic.3-Mic.12 population displayed a homeostatic microglial profile, the Act.1 subcluster showed robust expression of canonical DAM genes, supporting its classification as an AD-associated microglial state (Fig.4f). Notably, although expression of homeostatic microglial markers was comparable across genotypes, DAM-associated genes were expressed at lower levels in Act.1 microglia from AO mice than from APP/PS1 mice. Together with the reduced abundance of the Act.1 subcluster, these findings indicate that Ogg1 deficiency limits both the expansion and transcriptional activation of DAM during AD pathology.

To identify the microglial transcriptional changes prevented by Ogg1 deficiency during AD pathology, we focused on prevDEGs. Intersecting microglia-specific DEGs from the APP/PS1 vs WT and AO vs APP/PS1 comparison identified 129 prevDEGs, indicating broad reversal of AD associated microglial gene expression programs by loss of Ogg1 (Fig.4g). Notably, this set included regulators of microglial activation and immune signaling, including Cd83^34^, Cst7^35^, Lacc1^36^, and Clec7a^37^, all of which were expressed at lower levels in AO mice than in APP/PS1 mice (Fig.4h).

We next asked whether the Ogg1-dependent microglial transcriptional changes identified in APP/PS1 mice are conserved in human AD. To address this, we compared our microglial expression profiles with a published snRNA-seq dataset from the parietal cortex of individuals with AD^33^, including both sporadic AD (hAD) and autosomal-dominant AD (hADAD) cases carrying APP or PSEN1 mutations, which mirror the genetic alterations underlying our APP/PS1 mouse model (Fig.4i). Comparative analysis identified 34 genes shared between APP/PS1 and hADAD microglia, indicating that the APP/PS1 model recapitulates key aspects of human AD-associated microglial transcriptional changes. Notably, 41% of these genes were classified as prevDEGs, demonstrating that Ogg1 deficiency prevented dysregulation of a substantial fraction of conserved AD-associated transcriptional changes. Consistent with this finding, AO mice exhibited fewer significantly dysregulated genes overlapping with both hAD and hADAD. Together, these findings indicate that Ogg1 deficiency counteracts a subset of AD-associated microglial transcriptional changes conserved across mouse and human disease, suggesting the translational relevance of Ogg1-dependent microglial regulation.

### Loss of Ogg1 exerts context-dependent effects on astrocyte and oligodendrocyte transcriptional programs

Given the central role of astrocytes in maintaining synaptic function and neuronal homeostasis, processes disrupted in AD^38,39^, we next examined the astrocyte population. UMAP analysis resolved 15 transcriptionally distinct astrocyte subclusters. Functional annotation using GO enrichment, identified three subclusters (SynapticT Ast.1-3) enriched for synaptic transmission-related pathways (Fig.5a). Although overall cluster composition was comparable across genotypes (Supp.Fig9a), subcluster abundance differed substantially. SynapticT Ast.1 was significantly reduced in APP/PS1 relative to WT. In contrast, Ast.7, enriched for biological processes related to neuronal development and differentiation, were significantly increased in APP/PS1 compared to both WT and AO mice (Fig.5b). To determine whether these transcriptional changes were accompanied by astrocyte activation at the tissue level, we quantified GFAP (glial fibrillary acidic protein) immunoreactivity. APP/PS1 and AO mice exhibited significantly higher cortical GFAP immunoreactivity than WT, whereas hippocampal changes did not reach statistical significance (Supp.Fig9b and c). Consistent with the increased IBA1 immunoreactivity observed in *Ogg1^-/-^* mice, GFAP immunoreactivity was likewise elevated in both brain regions, suggesting that loss of Ogg1 promotes glial activation under physiological conditions (Fig.4c and d; Supp.Fig9b and c).

Astrocyte-specific differential expression analysis revealed a striking similarity between APP/PS1 and *Ogg1^-/-^* mice, which displayed comparable numbers of DEGs and a substantial overlap, particularly among downregulated genes (Supp.Fig9d). Shared downregulated genes were enriched for pathways related to synaptic transmission and synaptic signaling, with the strongest enrichment in *Ogg1^-/-^* mice (Fig.5c). Notably, these changes were absent in AO mice, indicating that Ogg1 deficiency induces astrocytic transcriptional programs linked to synaptic function under physiological conditions but not during amyloid pathology.

Consistent with these findings, we identified 302 astrocytic prevDEGs, indicating that Ogg1 deficiency markedly prevented transcriptional alterations observed in APP/PS1 astrocytes (Fig.5d). These genes included established regulators of neuroinflammation^40,41^ and synaptic plasticity^42^ (Fig.5e), as well as GFAP, whose increased expression in APP/PS1 mice was restored to WT-like levels in AO mice (Fig.5 e). Despite these transcriptional changes, hippocampal GFAP protein abundance remained unchanged, (Supp.Fig9b and c), suggesting that transcriptional remodeling precedes detectable protein-level changes at this stage of disease.

**Figure 5.**
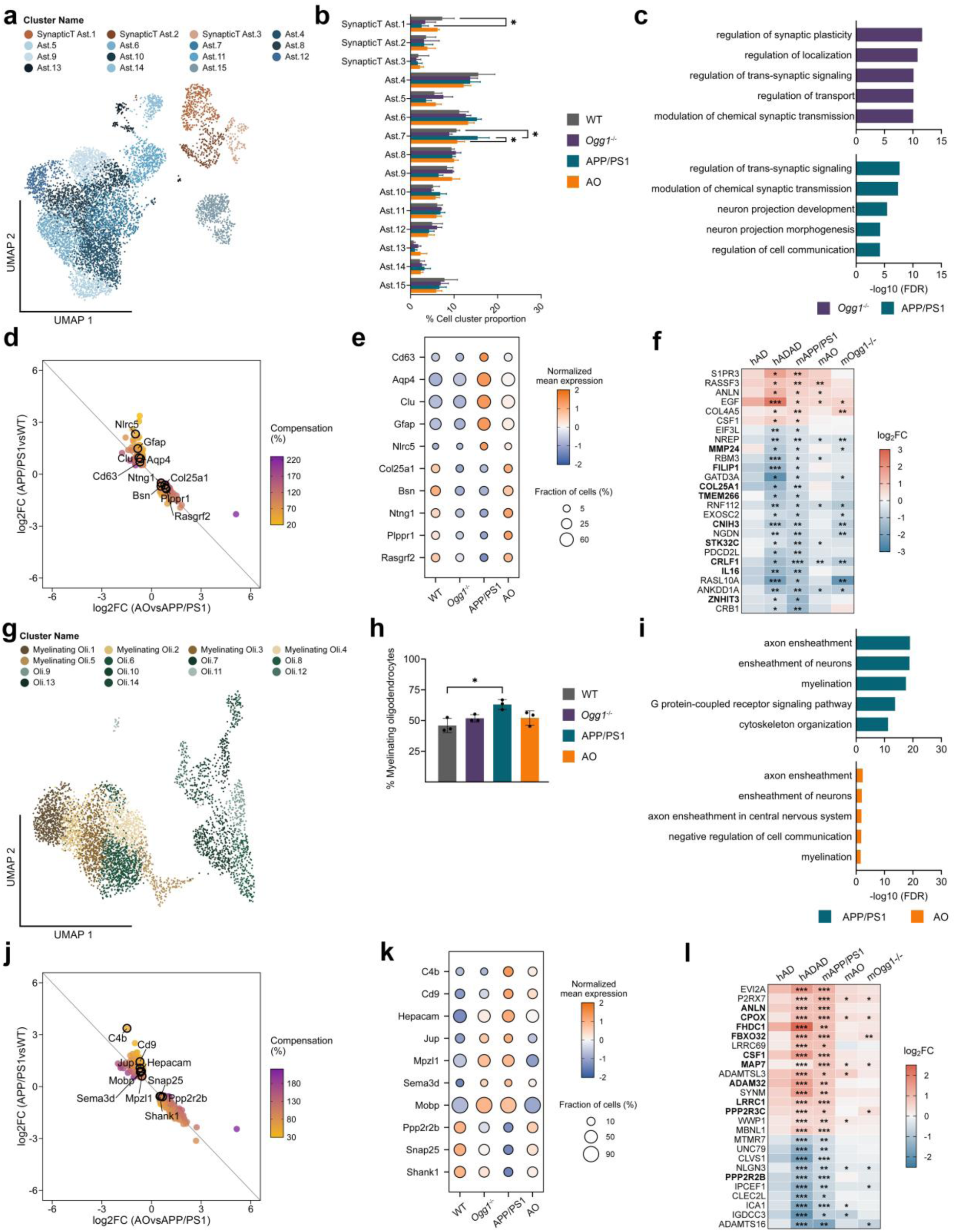
Loss of Ogg1 reshapes astrocyte and oligodendrocyte transcriptional programs in a context-dependent manner. (a) UMAP representation of astrocyte nuclei derived from snRNA-seq, revealing subclusters defined by transcriptomic similarity and functional annotation based on GO biological process enrichment. (b) Percentage distribution of nuclei among the astrocyte subclusters across all genotypes. Data are presented as mean ± SEM, *p < 0.05. (c) Gene ontology (GO) biological process enrichment of unique downregulated genes in the astrocyte cluster of *Ogg1^-/-^* and APP/PS1. (d) Scatter plot showing the relationship between gene expression changes in APP/PS1 versus WT and AO versus APP/PS1 astrocytes. Each dot represents a preventative differentially expressed gene (prevDEG). The color gradient indicates the compensation rate (%), calculated as the ratio of the log₂ fold changes in AO versus APP/PS1 to that in APP/PS1 versus WT, with higher values indicating greater prevention of AD-associated transcriptional dysregulation. Genes outlined in black indicate selected prevDEGs. (e) Dot plot depicting normalized mean expression of prevDEGs across genotypes in astrocytes. Dot size reflects the percentage of nuclei expressing the indicated gene. (f) Comparison of significantly DEGs (|log_2_ fold change| ≥ 0.5, p < 0.05) in astrocytes from our mouse snRNA-seq dataset and human parietal cortex astrocytes from sporadic AD (hAD) and autosomal dominant AD (hADAD; APP and PSEN1) reported by Brase et al.^33^. Shown are genes significantly dysregulated in hADAD (Benjamini–Hochberg-adjusted p < 0.05) with the same direction of change in APP/PS1 mice. Color indicates log₂ fold changes relative to WT mice (mouse) or neuropathology-free controls (human). Genes highlighted in bold represent prevDEGs. Asterisks indicate genes that are significantly differentially expressed (*p < 0.05, **p < 0.01, ***p < 0.001; Benjamini–Hochberg-adjusted p for human, nominal p for mouse comparisons). (g) UMAP representation of oligodendrocyte nuclei derived from snRNA-seq, revealing subclusters defined by transcriptomic similarity and functional annotation based on GO biological process enrichment. (h) Bar graph showing the relative proportion of myelinating oligodendrocytes across all genotypes. Data are presented as mean ± SEM, *p < 0.05. (i) GO biological process enrichment of unique upregulated genes in the oligodendrocyte cluster of APP/PS1 and AO. (j) Scatter plot showing the relationship between gene expression changes in APP/PS1 versus WT and AO versus APP/PS1 oligodendrocytes. Each dot represents a prevDEG. The color gradient indicates the compensation rate (%), calculated as the ratio of the log₂ fold changes in AO versus APP/PS1 to that in APP/PS1 versus WT, with higher values indicating greater prevention of AD-associated transcriptional dysregulation. Genes outlined in black indicate selected prevDEGs. (k) Dot plot depicting normalized mean expression of prevDEGs across genotypes in oligodendrocytes. Dot size reflects the percentage of nuclei expressing the indicated gene. (l) Comparison of significantly DEGs (|log_2_ fold change| ≥ 0.5, p < 0.05) in oligodendrocytes from our mouse snRNA-seq dataset and human parietal cortex oligodendrocytes from sporadic AD (hAD) and autosomal dominant AD (hADAD; APP and PSEN1) reported by Brase et al.^33^. Shown are genes significantly dysregulated in hADAD (Benjamini–Hochberg-adjusted p < 0.05) with the same direction of change in APP/PS1 mice. Color indicates log₂ fold changes relative to WT mice (mouse) or neuropathology-free controls (human). Genes highlighted in bold represent prevDEGs. Asterisks indicate genes that are significantly differentially expressed (*p < 0.05, **p < 0.01, ***p < 0.001; Benjamini–Hochberg-adjusted p for human, nominal p for mouse comparisons).

Recent studies have identified disease-associated astrocytes (DAAs) as a distinct astrocyte population that emerges during AD progression^43^. APP/PS1 astrocytes showed robust expression of canonical DAA genes, supporting the presence of a DAA-like transcriptional program in this model. Notably, expression of these genes was reduced in AO mice, indicating that Ogg1 deficiency prevents the establishment of DAA transcriptional programs during amyloid pathology (Supp.Fig9 e).

Comparison of astrocytic transcriptional profiles with the published AD snRNA-seq dataset^33^showed substantial overlap between APP/PS1 and hADAD astrocytes, which was reduced in AO (Fig.5f). Surprisingly, *Ogg1^-/-^* astrocytes also recapitulated many of the transcriptional alterations observed in human AD astrocytes, further supporting the notion that loss of Ogg1 in the absence of pathology induces AD-like astrocytic programs.

Given the essential role of oligodendrocytes in myelin maintenance and axonal support, and their emerging contribution to AD-associated circuit dysfunction^44^, we next examined the oligodendrocyte population. UMAP analysis resolved 14 transcriptionally distinct oligodendrocyte subclusters (Fig.5g). Functional annotation identified five subclusters (Myelinating Oli.1-5) enriched for myelination-related processes (Fig.5g). Although overall cluster composition was largely preserved across genotypes (Supp.Fig10a), selective abundance changes were observed. Oli.6 was increased in *Ogg1^-/-^* mice, whereas Oli.7 was reduced in APP/PS1 mice relative to WT, although neither cluster exhibited clear functional enrichment (Supp.Fig10b). In contrast, pooling all five myelinating oligodendrocyte subclusters revealed a significant increase in APP/PS1 mice that was absent in AO mice (Fig.5h).

Notably, oligodendrocytes were the most transcriptionally affected cell type in APP/PS1, with the highest number of DEGs (Supp.Fig10c). Upregulated genes were predominantly enriched for myelination-related pathways, whereas this enrichment was markedly attenuated in AO mice (Fig.5 i). As myelination is essential for axonal integrity and cognitive function^45^, the increased expression of these genes in APP/PS1 mice may reflect a compensatory response to disease-associated myelin dysfunction. Conversely, genes downregulated in APP/PS1 oligodendrocytes were strongly enriched for synaptic signaling pathways, an effect that was largely prevented by Ogg1 deficiency (Supp.Fig10 d). In *Ogg1^-/-^* mice, downregulated genes were instead primarily associated with developmental processes. Together, these findings suggest that loss of Ogg1 mitigates AD-associated remodeling of oligodendrocyte transcriptional programs.

To assess disease-associated oligodendrocyte states, we next examined a recently described disease-associated oligodendrocyte (DOL) signature identified across multiple neurodegenerative disorders^46^. APP/PS1 oligodendrocytes displayed increased expression of DOL-associated genes, (Supp.Fig10e), consistent with the activation of a disease-associated oligodendrocyte program. Notably, this response was reduced in AO mice, indicating that loss of Ogg1 attenuates the emergence of DOL-like transcriptional states during amyloid pathology.

Consistent with these observations, we identified 1,106 oligodendrocyte prevDEGs, representing the largest number of preventative transcriptional changes by Ogg1 among all examined cell types (Fig.5j). These genes include key regulators of myelin organization, such as Mobp^47^ and Mpzl1^48^, which were elevated in APP/PS1 mice but restored toward WT levels in AO mice (Fig.5k). Conversely, genes involved in synaptic function, including Shank1^49^ and the synaptic vesicle regulator Snap25^50^, were downregulated in APP/PS1 oligodendrocytes but restored by Ogg1 deficiency. Finally, comparison with human AD snRNAseq-data^33^ revealed a substantial overlap between APP/PS1 and human AD oligodendrocyte transcriptional signatures, whereas the overlap was considerably reduced in AO mice (Fig.5l). Similar to astrocytes, *Ogg1^-/-^* oligodendrocytes shared transcriptional features with human AD, indicating that Ogg1 deficiency alone is sufficient to induce aspects of an AD-like state (Fig.5k and l).

Together, these findings demonstrate that Ogg1 deficiency exerts context-dependent effects on astrocyte and oligodendrocyte transcriptional programs, promoting AD-like changes under physiological conditions while preventing disease-associated glia remodeling in the presence of amyloid pathology.

### Integrated transcriptomic and chromatin accessibility analyses reveal context-dependent glial remodeling in AD

To determine whether the transcriptional changes were accompanied by changes in promoter accessibility, we compared cell type-specific prevDEGs across bulk RNA-seq, ATAC-seq and snRNA-seq datasets. We focused on prevDEGs, as these represent transcriptional changes prevented in AO mice and therefore more closely reflect the protective phenotype. For each glial cell type, we examined the correlation between promoter accessibility and gene transcription and compared the effect sizes of transcriptional and chromatin changes across genotypes (Fig.6a).

**Figure 6.**
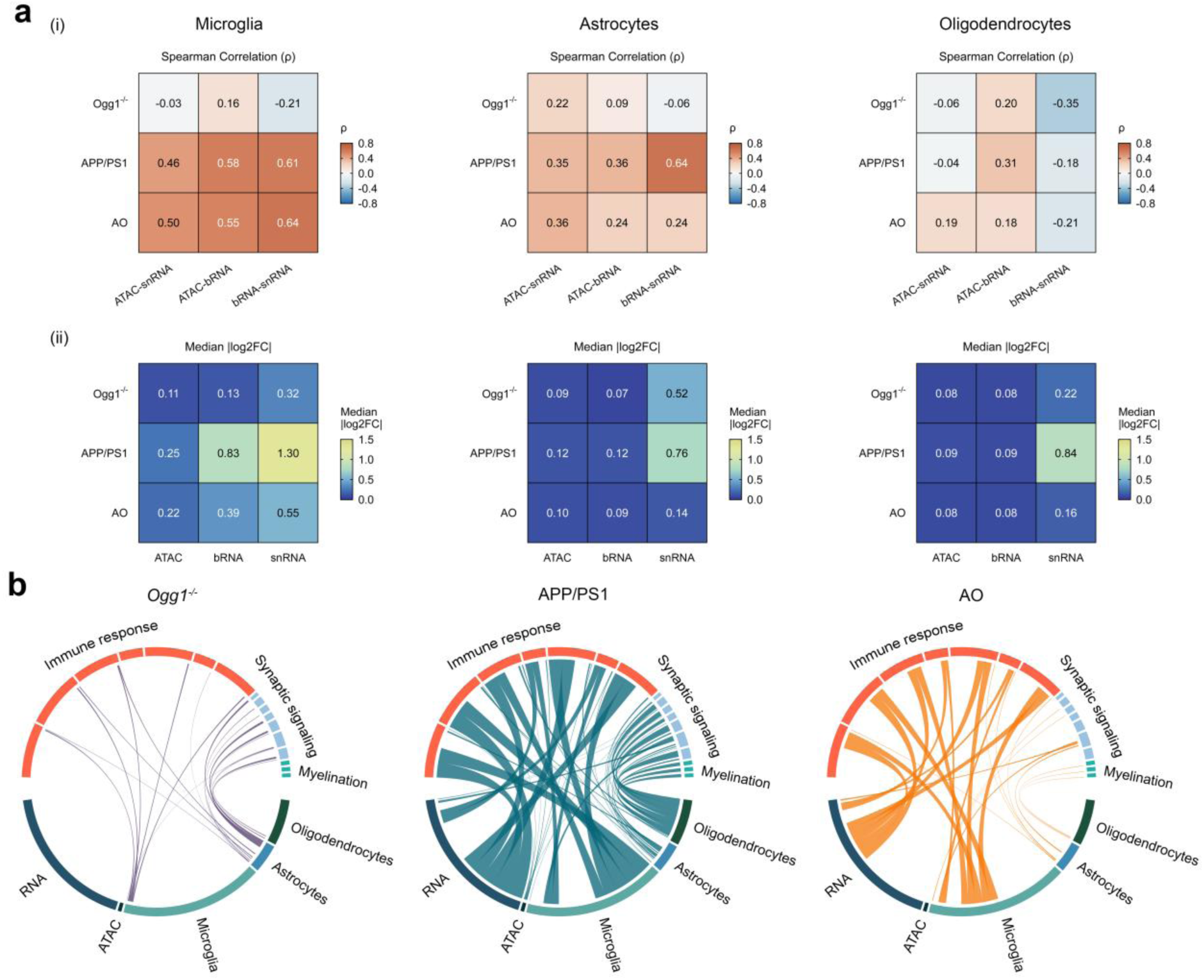
Integrated transcriptomic and chromatin accessibility analyses reveal Ogg1-dependent glial programs in AD. (a) Correlation between log₂ fold changes for cell type-specific preventative differentially expressed genes (prevDEGs) across promoter accessibility, bulk RNA-seq (bRNA), and snRNA-seq in microglia, astrocytes, and oligodendrocytes. Promoter accessibility was assessed by ATAC-seq, with log₂ fold changes calculated from promoter-annotated peaks and averaged when multiple peaks mapped to the same promoter. (i) Spearman’s rank correlation coefficient (ρ) describing the association between log₂ fold changes for each assay pair, calculated separately for each cell type and genotype. (ii) Median absolute log₂ fold change (|log₂FC|) summarizing the effect size for each assay and genotype, irrespective of direction. (b) Circular chord diagrams illustrating convergence of gene ontology (GO) biological process enrichment across bRNA, ATAC-seq, and cell type-specific snRNA-seq datasets (microglia, astrocytes, and oligodendrocytes) in *Ogg1^-/-^*, APP/PS1, and AO mice. Chords connect significantly enriched GO terms shared between datasets, with chord thickness proportional to pathway significance (-log10 FDR). Related GO terms are organized into major biological processes, including immune response, synaptic signaling, and myelination.

In APP/PS1 mice, prevDEGs showed moderate correlations between promoter accessibility and gene transcription in microglia and astrocytes, but weaker correlations in oligodendrocytes. A similar pattern was observed in AO mice, indicating that these correlations were largely preserved, whereas correlations in *Ogg1^-/-^* mice were generally weak or negative (Fig.6a i).

Although correlations between promoter accessibility and gene transcription were similar across genotypes, consistent with epigenetic regulation of prevDEGs, the magnitude of the associated transcriptional and chromatin changes differed substantially (Fig.6a ii). Across all glial cell types, snRNA-seq showed the largest median absolute log₂ fold changes with effect sizes being greatest in APP/PS1 mice, intermediate in *Ogg1^-/-^* mice, and substantially reduced in AO mice. Notably, the effect size in astrocytes was similar in *Ogg1^-/-^* and APP/PS1 mice but markedly reduced in AO mice. Consistent with this, both bRNA-seq and ATAC-seq showed reduced effect sizes in AO mice, particularly in microglia.

To determine whether these coordinated molecular changes converged on common biological pathways, we integrated enrichment analyses from bRNA-seq, ATAC-seq, and cell type-specific snRNA-seq datasets. This analysis revealed distinct genotype-dependent patterns across molecular layers (Fig.6b). *Ogg1^-/-^* mice exhibited a limited set of pathway associations, some of which partially overlapped with those observed in AD. In contrast, APP/PS1 mice showed widespread enrichment of immune-, synaptic-, and myelination-related pathways across the integrated datasets, consistent with extensive transcriptional and chromatin remodeling in AD. These pathway associations were markedly reduced in number and strength in AO mice, consistent with a broad attenuation of AD-associated molecular remodeling following Ogg1 deficiency. Together, these findings identify Ogg1 as a context-dependent regulator of glial transcriptional and chromatin remodeling in AD.

## Discussion

While Ogg1 is best known for its role in repairing oxidative DNA lesions, our findings identify a previously unrecognized role for Ogg1 in regulating transcriptional programs and cognitive function in AD. By integrating behavioral, transcriptional, and chromatin accessibility analyses, we show that loss of Ogg1 alters neuronal and glial responses and partially prevents AD-associated pathology. Together, these findings suggest that Ogg1 contributes to AD through mechanisms extending beyond its canonical role in BER.

### Ogg1 is a context-dependent regulator of AD pathology

A major finding of our study is that the biological consequences of Ogg1 deficiency are context dependent. In the healthy brain, Ogg1 may support glial homeostasis, such that its loss disrupts gene expression required for homeostatic glial function. Consistent with this, maintenance of glial homeostasis depends on active gene expression, and even modest transcriptional changes can drive inflammatory state transitions in the absence of overt pathology^51^. In contrast, in the presence of amyloid pathology loss of Ogg1 appears to limit AD-associated glial activation and the accompanying transcriptional changes. This context dependence may also account for differences with previous studies. The memory impairment observed in Ogg1-deficient mice differs from previous work^52^, showing no cognitive deficits following *Ogg1* deletion. One possible explanation is the use of different C57BL/6 substrains, which vary in hippocampal function, immune responses, and behavior^53^. Thus, genetic background may contribute to the context-dependent effects of Ogg1 loss.

Our findings also differ from previous reports showing accelerated oxidative damage and microgliosis following Ogg1 deficiency in the AppNL-G-F knock-in model of AD^54^. Differences in amyloid pathology, immune responses, and disease progression between APP/PS1 and AppNL-G-F models may contribute to these distinct outcomes. In addition, the previous study was conducted in female mice, whereas our analyses were performed in male mice. As biological sex influences AD pathophysiology and neuroimmune responses^55^, it may represent an additional factor contributing to the context-dependent effects of Ogg1 loss. Further support for context-dependent Ogg1 function comes from studies showing that increased mitochondrial OGG1 expression reduces age-associated neuroinflammation^56^, whereas pharmacological inhibition of OGG1 suppresses NF-κB-dependent inflammatory gene expression^57^.

### Ogg1 mediates AD pathology through transcriptional and epigenetic regulation rather than canonical DNA repair

The absence of detectable differences in 8-oxo-dG levels between genotypes suggests that defective oxidative lesion repair is unlikely to be the primary mechanism underlying the observed phenotypes. Although global measurements cannot exclude locus-specific effects at regulatory elements, our findings support a role for Ogg1 in transcriptional regulation through chromatin-associated mechanisms in AD. This is consistent with evidence that Ogg1 contributes to gene regulation beyond its canonical DNA repair function, including through interactions with chromatin-associated regulators^52,58^. Ogg1 can promote transcription through recruitment of transcription factors and epigenetic regulators^59^, while the DNA repair deficient OGG1S326C variant exhibits prolonged DNA binding and promotes inflammatory gene expression^60^. Consistent with this, work from our group demonstrated that genome-wide 8-oxoG distribution and OGG1-dependent transcriptional changes are largely uncoupled^61^, suggesting that oxidative lesion accumulation and OGG1-mediated gene regulation may operate through partly independent mechanisms.

Several of our observations suggest that Ogg1 influences chromatin organization in AD. Many genes dysregulated in APP/PS1 mice also exhibited altered chromatin accessibility, and both transcriptional and accessibility changes were partially prevented by Ogg1 deficiency. Notably, promoter accessibility of prevDEGs remained associated with changes in gene expression in AO mice, albeit with substantially reduced effect sizes. In APP/PS1 mice, regions uniquely altered in accessibility were enriched for motifs linked to immune signaling, particularly PU.1^62^. Prevention of these accessibility changes in AO mice implicates Ogg1 in transcription factor networks regulating microglial activation, potentially involving PU.1, as supported by our findings and previous studies^63,64^. However, whether this reflects altered PU.1 occupancy or chromatin accessibility remains unclear.

The effects of Ogg1 deficiency were also evident at loci involved in higher-order genome organization, including the Firre locus, where it prevented the APP/PS1-associated increase in chromatin accessibility. Firre coordinates chromatin organization and long-range genomic interactions^26^, raising the possibility that Ogg1-dependent regulation at this locus contributes to broader chromatin remodeling during AD.

Furthermore, Ogg1 may influence local chromatin accessibility and transcription through interactions with oxidized regulatory regions and recruitment of transcriptional and epigenetic regulators^59^. Although global 8-oxo-dG levels were unchanged in our study, locus-specific interactions between Ogg1 and oxidative DNA lesions cannot be excluded. Oxidative lesions at guanine-rich regions may alter local DNA topology, including G-quadruplex formation, with potential consequences for transcriptional regulation^65,66^. Alternatively, Ogg1 may influence chromatin accessibility through non-canonical mechanisms beyond oxidative DNA repair. This possibility is consistent with our previous work demonstrating that Ogg1 regulates PRC2 occupancy and Polycomb-associated chromatin states^52^. Whether Ogg1-dependent chromatin remodeling in AD involves recognition of 8-oxo-dG, recruitment of epigenetic regulators, or both remains to be determined. Nevertheless, our findings support a non-canonical role for Ogg1 in chromatin regulation and identify chromatin remodeling as a potential mechanism through which Ogg1 shapes AD-associated transcriptional responses.

### Ogg1 deficiency prevents AD-associated transcriptional dysregulation across glial cell types

The protective effects of Ogg1 deficiency were most pronounced across glial populations, where loss of Ogg1 broadly attenuated AD-associated transcriptional dysregulation. This is consistent with evidence that glial dysfunction contributes to AD progression^67^.

Microglia exhibited extensive transcriptional changes following Ogg1 deficiency, consistent with their established role in amyloid pathology and neuroinflammation in AD^68^. Loss of Ogg1 reduced the emergence of a disease-associated Act1 microglial cluster resembling DAM states^35^, prevented AD-associated inflammatory transcriptional changes, and promoted a more homeostatic microglial profile accompanied by reduced activation. These changes may contribute to the reduced insoluble Aβ accumulation observed in AO mice, as inflammatory microglial states can impair Aβ clearance through NLRP3-dependent pathways^69^. Notably, insoluble Aβ was reduced without changes in overall plaque burden, suggesting that Ogg1 deficiency may not primarily affect amyloid production but instead alter how existing plaques are processed by microglia, potentially through changes in plaque remodeling and compaction^70^.

Astrocyte and oligodendrocytes also underwent substantial Ogg1-dependent transcriptional remodeling. Notably, Ogg1 deficiency alone induced aspects of an AD-like transcriptional state in both cell types, whereas these changes were attenuated in APP/PS1 mice lacking Ogg1. In astrocytes, the affected genes were enriched for synaptic signaling pathways, which may contribute to the memory deficits observed in *Ogg1^-/-^* mice through impaired regulation of synaptic function and plasticity^71^. In oligodendrocytes, APP/PS1 mice showed dysregulation of myelination-related pathways which are among the earliest plaque-associated transcriptional responses and preceding inflammatory gene programs that emerge at later disease stages^72^. Upregulation of myelination-related genes may therefore reflect a compensatory adaptation to neuronal dysfunction.

### Ogg1-dependent glial programs are conserved in human AD

APP/PS1-associated transcriptional changes overlapped substantially with human AD signatures across glial populations, despite recognized differences between mouse models and human disease. Microglia showed the strongest overlap with human AD signatures, suggesting that Ogg1-dependent microglial transcriptional changes in experimental AD models are also relevant to human disease^73^.

Additional support for the relevance of OGG1 in human AD comes from genetic and postmortem studies. *OGG1* polymorphisms associated with AD alter enzymatic activity and cellular responses to oxidative stress^11^. Increased OGG1 expression has been reported in individuals with preclinical AD and in patients with mild cognitive impairment (MCI)^74,75^. Consistent with these findings, we observed elevated OGG1 expression at both the transcription and protein levels in patients with AD. Notably, despite this increase in expression, OGG1 glycosylase activity is reduced during MCI and late-stage AD^75^, suggesting that transcriptional upregulation of OGG1 may represent a compensatory response to impaired oxidative DNA repair. Together, these findings support the relevance of OGG1-dependent mechanisms in human AD.

### Limitations and conclusion

Several limitations should be considered when interpreting these findings. The use of a constitutive *Ogg1* knockout model does not distinguish between developmental and disease stage-specific effects. In addition, analyses at a single disease stage and bulk epigenomic profiling limit the resolution of temporal and cell type-specific mechanisms. Although APP/PS1 mice recapitulate key aspects of amyloid pathology, they do not model tau pathology or the extent of neurodegeneration observed in human AD. Future studies using inducible and cell type-specific approaches, higher-resolution epigenomic analyses, and human cellular systems will help clarify how Ogg1 regulates chromatin and transcription during AD progression and determine its potential as a therapeutic target.

In conclusion, our findings identify Ogg1 as a context-dependent regulator of glial states and reveal a role in transcriptional and chromatin remodeling in AD. These findings expand the known functions of Ogg1 beyond oxidative DNA repair and identify Ogg1 as a previously unrecognized contributor to AD pathogenesis.

## Materials and Methods

### Animals

All experiments were approved by the Norwegian Animal Research Authority (NARA) and conducted in accordance with the laws and regulations controlling experimental procedures in live animals in Norway and the European Union’s Directive 86/609/EEC. Mice were housed under standard laboratory conditions of 19-22°C at the Comparative Medicine Core Facility (Trondheim, Norway) in a 12 h reversed light/dark cycle with access to food and water *ad libitum*. All mice used in this study were maintained on a C57BL/6J background. APP/PS1 mice previously described^76^ which were generally bred heterozygous and *Ogg1^-/-^* mice previously described^77^ were used to generate APP/PS1 x *Ogg1^-/-^* (AO) mice. APP/PS1 transgenic mice overexpress a human APP with Swedish mutation (KM670/671NL) and PS1 with L166P mutation under the control of the neuron specific promoter Thy1^76^. C57BL/6J wild-type (WT) mice, not carrying human APP/PS1 genes, were littermates of APP/PS1 mice and were used as control groups. For all experiments 6 months old male mice were used. Mouse litters were genotyped using genomic DNA obtained from ear biopsies or tail biopsies after animals were euthanized.

### Mouse genomic DNA isolation

Genomic DNA was isolated from mouse hippocampal and cortical tissue using the DNeasy Blood C Tissue Kit (Qiagen) according to the manufacturer’s instructions. To minimize artifactual DNA oxidation during isolation, butylated hydroxytoluene (BHT) and deferoxamine (DFO) were included in all buffers at final concentrations of 0.1 mM each. DNA concentrations were determined using a NanoDrop spectrophotometer (Thermo Fisher Scientific).

### Global 8-oxo-dG quantification

Global 8-oxo-2′-deoxyguanosine (8-oxo-dG) levels were quantified in genomic DNA isolated from hippocampal and cortical tissue of WT, APP/PS1, *Ogg1^-/-^* and AO mice. Genomic DNA was purified by isopropanol precipitation using ammonium acetate, washed twice with 70% ethanol, air-dried, and resuspended in nuclease-free water. DNA was enzymatically hydrolyzed for 1 h at 37 °C in hydrolysis buffer (10 mM NH₄Ac, 1 mM MgCl₂, 0.1 mM ZnCl₂) containing nuclease P1 (0.008 U/μl), benzonase (0.8 U/μl), alkaline phosphatase (0.04 U/μl), and the antioxidants butylated hydroxytoluene (BHT) and chelator deferoxamine (DFO) (0.1 mM each). Proteins were precipitated by addition of ice-cold acetonitrile followed by centrifugation (16,000 × g, 30 min, 4 °C). Supernatants were lyophilized overnight at −80 °C and resuspended in water.

8-oxo-dG was quantified by liquid chromatography-tandem mass spectrometry (LC-MS/MS) using an Agilent 6495 Triple Quadrupole LC/MS system equipped with an Eclipse Plus C18 RRHD column (2.1 × 150 mm, 1.8 μm). The 8-oxo-dG transition monitored was m/z 284.1 → 168.0. Unmodified deoxyribonucleosides were quantified separately using an API5000 Triple Quadrupole mass spectrometer (Applied Biosystems) and an Acentis Express C18 column (0.5 × 150 mm, 2.7 μm) to normalize 8-oxo-dG levels. Chromatographic conditions and mass transitions for canonical nucleosides (dA, dC, dG, and dT) were as previously described^61^.

### Human Aβ42 and Aβ40 measurement

Enzyme-linked immunosorbent assay (ELISA) was conducted by using Human Aß40 ELISA Kit and Human Aß42 ELISA Kit (Thermo Fisher Scientific). In brief, entire mouse brain samples were homogenized in carbonate buffer (pH 11.5, 100 mM Na_2_CO_3_, 50 mM NaCl, protease inhibitor) using a MagNA Lyser (Roche) for 5 s at 5000 speed followed by a centrifugation step at 16400 rpm for 90 min. The supernatant was collected and diluted in 8 M guanidine buffer (pH 8.0, guanidine/HCl 8.2 M, Tris/HCl 82 mM) to obtain the soluble Aβ fractions. For the insoluble Aβ fractions, the remaining pellets were resuspended in 5 M guanidine buffer (pH 8, guanidine/HCl 5 M, Tris/HCl 50 mM) and shaken at room temperature for 3 h followed by a centrifugation step at 1400 rpm for 20 min. Protein concentrations were estimated with a ND-1000 spectrophotometer (Nanodrop Technologies). Aβ42 and Aβ40 concentrations were measured with the Human Aβ42 and Human Aβ40 ELISA kits (Thermo Fisher), following the manufacturer’s protocol. Absorbance was read at 450 nm using a FLUOstar Omega microplate reader (BMG Labtech).

### Immunohistochemistry

Mice were transcardially perfused with PBS, and brains were dissected, fixed in 4% paraformaldehyde for at least 48 h, paraffin-embedded, and cut into 4-μm coronal sections. Sections were deparaffinized, rehydrated, and subjected to antigen retrieval in citrate buffer (pH 6.0) at 100°C under pressure. Immunohistochemistry was performed using the Mouse and Rabbit Specific HRP/DAB (ABC) Detection IHC Kit (Abcam) and primary antibodies against Amyloid β (Aβ) (6F/3D, Thermo Fisher, 1:100), GFAP (Z0334, Dako, 1:2,000), and IBA1 (019-19741, Fujifilm Wako, 1:1,000). Sections were incubated with primary antibodies for 1 h at room temperature. Staining was visualized using streptavidin-peroxidase and DAB, followed by hematoxylin and eosin counterstaining. Finally, sections were dehydrated and mounted using Entellan (Sigma-Aldrich).

### Image quantification

Stained sections were scanned by the Cellular and Molecular Imaging Core facility (CMIC, NTNU) using an Olympus VS120S5 with a 20× objective, on a single Z-plane with low focus grid density. Plaque quantification, as well as the abundance of astrocytes and microglia, were analyzed in ImageJ by measuring the area of interest and analyzing the particles using a specified threshold and minimum particle size. Microglial activation was evaluated as previously described^31^. In short, total microglia cells and cell bodies were measured independently. Finally, the cell body to cell size ratio was calculated and used as an indicator for microglia activation. For all quantifications, the average of at least two sections per animal stained in independent runs were used.

### Open Field

Exploratory behavior, anxiety-like behavior, and general locomotor activity were examined using the open-field test as previously described^78^. Prior to the open-field test, mice were handled 5 min for five consecutive days and mice placed individually into an open-field apparatus (50 x 50 x 30 cm open field area), which was virtually divided into 3 × 3 subareas as one center area, four corner areas, and four side areas. Each mouse was kept in the open-field apparatus for ten min while the total locomotor activity (time spent in different areas and distance traveled) was recorded automatically and analyzed with the Any-Maze tracking software (Stoelting Europe).

### Novel object location

After mice were subjected to the open-field test, animals were habituated to the open-field apparatus for 5 min for two additional consecutive days in the absence of objects, as previously described^52^. During training, mice were placed into the experimental apparatus containing two identical objects (Duplo blocks) and allowed to explore for 10 min. Short-term and long-term retention tests were conducted by placing the mice back into the experimental apparatus for 10 min, either 1 h or 24 h after training. To assess spatial object location-dependent memory, the placement of one familiar object remained unchanged whereas the other familiar object was moved and placed adjacent to a different corner inside the experimental apparatus. Exploration was scored when the head of the animal entered the virtual zone adjacent to the object. All training and testing trials were videotaped and analyzed with Any-Maze software (Stoelting Europe). The relative exploration time (t) was expressed as a percent discrimination index (D.I. = t_novel_ / (t_novel_ + t_familiar_) × 100). Mean exploration times were then calculated and the discrimination indices between experimental groups were compared. Animals that explored each object for less than 6 seconds during training or explored both objects for less than 10 seconds total during testing were excluded from the analysis.

### Tissue collection

Animals were sacrificed by cervical dislocation, and hippocampi were dissected, isolated, snap-frozen in liquid nitrogen and stored at −80°C until further use. All mice included in high throughput sequencing experiments were re-genotyped by using genomic DNA obtained from tail biopsies after being sacrificed.

### Bulk RNA-sequencing

Snap-frozen hippocampi from two animals of the same genotype were combined and dissociated in 1 ml of cold homogenization buffer following the manufacturer’s instructions (Diagenode ATAC-seq kit, Cat. No. C01080006). In brief, hippocampal tissue was gently homogenized by applying pestle “loose” and “tight” 40 times, respectively. All buffers during the tissue lysis contained 20 U/ml of RNase Inhibitor (Applied Biosystems, N8080119). Three independent biological replicates per genotype were performed. One part of the cell suspension was stored in 1500 μl Trizol (Invitrogen, 15596026) at −80°C for later processing, while the remaining part was utilized for nuclei isolation for ATAC-seq. For RNA extraction the Trizol samples were thawn on ice. After adding 400 μl chloroform, the samples were mixed by vortexing and then centrifuged at 14000 rpm for 15 min at 4°C. Subsequently, the clear aqueous layer was collected, and an equal volume of 100% ethanol was added and mixed. RNA was then extracted and isolated using the Qiagen RNeasy Kit (Qiagen, 74104), including DNA digestion as per the manufacturer’s protocol. RNA concentration and quality was determined using a Qubit HS RNA Assay Kit (Thermo Fisher, Q32852) and an Agilent 2100 Bioanalyzer, respectively. Sequencing (paired-end, 100 bp) was performed at BGI Tech Solutions Co., Hong Kong using a DNBSEQ-T7 sequencer with an average depth of 35 million reads per sample.

### ATAC-sequencing

Because ATAC-seq was paired with RNA-seq, the same cell suspension was used to isolate nuclei. Nuclei were collected after an iodixanol-based density gradient centrifugation and ATAC-seq was performed according to manufacturer’s instructions (Diagenode ATAC-seq kit, C01080006). Three independent biological replicates per genotype were performed. In brief, 50.000 hippocampal nuclei per replicate from all genotypes underwent tagmentation for 30 min at 37 °C. The tagmented DNA was then purified and amplified through PCR amplification for library generation using unique dual indexes (UDI). Size selection was conducted by using AMPure XP beads (Beckman Coulter, A63880) and DNA fragment size was determined by an Agilent 2100 Bioanalyzer. Concentration of the final libraries was measured with a Qubit dsDNA HS Assay Kit (Thermo Fisher, Q32851) and was further validated by qPCR using a KAPA Library Quantification Kit (Roche, KK4824) before being sequencing. Sequencing (paired-end, 100 bp) was performed at BGI Tech Solutions Co., Hong Kong using a DNBSEQ-G400 sequencer with an average depth of 28 million reads per sample.

### Single-nuclei RNA-sequencing

Frozen hippocampal tissue was homogenized and nuclei were isolated as previously described, with some minor modifications^52^. In brief, for each replicate snap-frozen hippocampi from 3 animals of the same genotype were pooled and homogenized by using Dounce homogenizers in 1 ml low sucrose buffer (0.32 M sucrose, 10 mM HEPES pH 8.0, 5 mM CaCl_2_, 3 mM Mg(CH_3_COO)_2_, 0.1 mM EDTA, 0.1% Triton X-100, 1 mM DTT) containing 20 U/ml of RNase Inhibitor (Applied Biosystems, N8080119). All steps were performed at 4°C or on ice. The homogenate was then pelleted by centrifugation at 350 g for 5 min at 4 °C, re-suspended in 1 ml of low sucrose buffer containing 20 U/ml of RNase Inhibitor and filtered through a 40 μm strainer. The volume of the cell suspension was then adjusted to 1.5 ml with low sucrose buffer containing 20 U/ml of RNase Inhibitor and extracted nuclei were purified through 6 ml sucrose cushion (10 mM HEPES pH 8.0, 1 M sucrose, 3 mM Mg(CH_3_COO)_2_, 1 mM DTT) by centrifugation at 3200 g for 10 min. Afterwards, purified nuclei were re-suspended in 1x PBS containing 1% BSA and 20 U/ml of RNase Inhibitor and the number of nuclei were counted with a hemocytometer. Nuclear fixation was performed for 3 million nuclei per sample according to the manufacturer’s instructions (Evercode Nuclei Fixation Kit v2, Parse Biosciences). After nuclei fixation, samples were cryopreserved with DMSO and stored in aliquots at -80 °C until the commencement of single nuclei RNA-Seq.

SnRNA-Seq was performed according to the manufacturer’s instructions (Evercode WT v2 Kit, Parse Biosciences). Briefly, RNA from fixed and permeabilized nuclei underwent reverse transcription and multiple rounds of ligation barcoding in customized 96-well plates, that contained uniquely barcoded primers in each well. Barcoding was performed simultaneously on all three independent biological replicates of each genotype, using the same number of nuclei as input for each sample. After three rounds of barcoding, nuclei were pooled together and equally distributed into 8 sublibraries. Nuclei were then lysed, and the barcoded cDNA was extracted and prepared for sequencing following the manufacturer’s instructions. In brief, barcoded cDNA from each sublibrary was amplified and then subjected to fragmentation, end repair and A-tailing followed by a double sided SPRI size selection and adaptor ligation. Finally, a specific UDI for each sublibrary was introduced by PCR and a double sided SPRI size selection was performed. Concentration of the final libraries was measured with a Qubit dsDNA HS Assay Kit (Thermo Fisher, Q32851) and cDNA fragment size was determined by an Agilent 2100 Bioanalyzer. Sublibraries were then pooled and sequencing (paired-end, 150 bp) was performed at the Norwegian Sequencing Centre (Oslo) using an Illumina Novaseq 6000 S4 platform with a sequencing depth of 50.000 read pairs/nuclei.

### Genomic data processing Bulk RNA-seq data

Adaptors were trimmed from reads using Cutadapt (v 2.8) and aligned to the GRCm38 genome using Hisat2 (v. 2.2.1). Properly mapped and paired reads were sorted and indexed using samtools (v. 1.10). For bigwigs visualization replicates were merged using samtools and bigwigs were generated using Deeptools. PCA was generated using Deeptools multiBAMsummary. Reads were counted using featureCounts from subread (v. 2.0.0) over protein-coding genes extracted from the Ensembl GRCm38 gtf (v. 102). Differential enrichment was determined using the quasilikelihood F-test in edgeR (v. 3.42.4) and genes were considered significant if they had a p value < 0.05 and the absolute log_2_ fold change was ≥ 0.5. Venn diagrams were created using Venny (v. 2.1).

### ATAC-seq data

Adaptors were trimmed from reads using Cutadapt (v 2.8) and aligned to the GRCm38 genome using Bowtie2 (v. 2.3.5.1). Duplicates were marked and reads were sorted and indexed using samtools (v. 1.10). For bigwigs visualization replicates were merged using samtools and bigwigs were generated using Deeptools normalized by RPKM. PCA was generated using Deeptools multiBAMsummary. Peaks were called using MACS3 (v. 3.0.0a7) narrowPeaks with the following settings: --nomodel -f BAMPE -q .05. Resulting peaks were combined and flattened using bedtools. Final peaks were annotated using ChIPseeker (v. 1.36.0) and were converted to SAF in R (v. 4.3.2). Reads were counted using featureCounts from subread (v. 2.0.0). Differential enrichment was determined using the quasilikelihood F-test in edgeR (v. 3.42.4) where peaks were considered significantly differentially enriched if the p value was < 0.05 and the absolute log_2_ fold change was ≥ 1. Venn diagrams were created using Venny (v. 2.1) by gene name. Differentially enriched regions were exported as bed files from R and motif analysis was done using Homer (v. 5.1). Gene tracks were generated in R using the Bioconductor package rtracklayer^79^.

### SnRNA-seq data

SnRNA-seq data were processed using Trailmaker™ (Parse Biosciences). Low-quality barcodes were removed using sample-specific minimum transcript thresholds determined from the distribution of transcripts per nucleus (870-1,650 transcripts). Nuclei with > 0.5% mitochondrial transcripts, outliers in the relationship between transcript counts and detected genes (identified by spline regression; sample-specific P = 1.2 × 10⁻⁴-3.4 × 10⁻⁴), and predicted doublets (scDblFinder probability threshold 0.77-0.86) were excluded. High-quality nuclei were normalized using SCTransform and integrated by reciprocal PCA (27 principal components) based on the 2,000 most variable genes after exclusion of ribosomal and mitochondrial genes. Clusters were identified using the Louvain algorithm (resolution = 0.8) and visualized by UMAP using a cosine distance metric (minimum distance = 0.3). Cluster marker genes were identified using the presto implementation of the Wilcoxon rank-sum test. Cell types were annotated using ScType, a marker-based cell-type identification method^80^. Genes were considered significant at p < 0.05 and absolute log₂ fold change ≥ 0.5. Venn diagrams were created using Venny (v2.1).

### Subclustering

Each annotated cell type was subclustered to resolve transcriptionally distinct populations. Nuclei from one cell type at a time were extracted and reprocessed in Trailmaker using the same workflow and parameters as for global clustering (SCTransform normalization, 2,000 recalculated highly variable genes excluding ribosomal and mitochondrial genes, RPCA integration with 27 principal components, Louvain clustering at resolution 0.8, and UMAP with cosine metric and minimum distance 0.3).

### Comparison with published disease-associated signatures

Published signatures of disease-associated microglia (DAM; Keren-Shaul et al.^35^), disease-associated astrocytes (DAA; Habib et al.^43^) and disease-associated oligodendrocytes (DOL; Kenigsbuch et al.^46^) were compared with the differential expression results of the corresponding cell types. For microglia, up- and downregulated DAM genes were taken from Supplementary Table 3 of Keren-Shaul et al. (Mann-Whitney p < 0.05, |log₂FC| > 1 relative to homeostatic microglia) and intersected with genes up- or downregulated in the activated microglia subcluster Act.1 relative to the non-activated microglia subclusters (excluding the second activated subcluster, Act.2) in APP/PS1 mice, retaining only genes with the same direction of change in both comparisons. For astrocytes, DAA genes were defined as genes differentially expressed between DAA (cluster 4) and GFAP-low homeostatic astrocytes (cluster 1) in Supplementary Table 2 of Habib et al.) and intersected with genes up- or downregulated in APP/PS1 astrocytes, retaining genes with the same direction of change in both datasets. For oligodendrocytes, the DOL gene list was obtained from the author’s repository (https://github.com/PierreBSC/DOL_project) and filtered for genes upregulated in APP/PS1 oligodendrocytes.

The resulting gene sets were visualized as heatmaps in R using ComplexHeatmap^81^. For microglia, log-normalized expression values were extracted from the Seurat object, back-transformed to the linear scale, and averaged per gene for Act.1 (APP/PS1 and AO mice; Act.1 was absent in WT and Ogg1 mice) and for the pooled non-activated microglia subclusters of each genotype (WT, *Ogg1^-/-^*, APP/PS1, AO). Mean expression was z-scored per gene across these six groups. For astrocytes and oligodendrocytes, mean expression per genotype was exported from Trailmaker and plotted without scaling. Genes were ordered by hierarchical clustering of the plotted values.

### Gene ontology enrichment analysis

Gene ontology (GO) enrichment was performed with the PANTHER overrepresentation test^82^ using the “GO biological process complete” annotation set and Fisher’s exact test with false discovery rate (FDR) correction. For bulk RNA-seq and snRNA-seq, up- and downregulated genes were analysed separately against all Mus musculus genes as reference list. For ATAC-seq, genes associated with differentially accessible regions were analysed against a custom reference list of all genes associated with consensus peaks. Terms with FDR < 0.05 were considered significantly enriched. GO enrichment graphs were generated in GraphPad Prism (v10; GraphPad Software, La Jolla, CA, USA).

### Preventative analysis

Preventative differentially expressed genes (prevDEGs) and differentially accessible regions (prevDARs) were identified by intersecting significant differential expression or accessibility results from the APP/PS1 versus WT and APP/PS1 versus AO comparisons. Only genes or regions that were significantly altered in both comparisons were retained. The degree of compensation was defined as [log_2_ fold change (APP/PS1 vs AO) / log_2_ fold change (APP/PS1 vs WT)]. The same approach was applied to bulk RNA-seq, ATAC-seq, and snRNA-seq data, with compensation rates calculated separately for each dataset or annotated cell type. A complete list of prevDEGs and prevDARs is provided in Supplementary Table 1.

### Dot plots of prevDEGs and prevDARs

Dot plots of selected prevDEGs and prevDARs were generated in R using ggplot2^83^. For bulk RNA-seq, mean FPKM per genotype was z-scored per gene across the four genotypes. For ATAC-seq, mean normalized chromatin accessibility per genotype was averaged across peaks annotated to the same gene and z-scored per gene across genotypes. For snRNA-seq, scaled mean expression (z-score across genotypes) and the percentage of nuclei expressing each gene were exported per genotype from Trailmaker’s dot plot function for the respective cell type. In all plots, dot colour represents the z-scored value; for snRNA-seq, dot size additionally represents the percentage of expressing nuclei. Genes were ordered by their value in APP/PS1 mice.

### Comparison with human data

For comparison with human AD, we used the snRNA-seq dataset of Brase et al.^33^, comprising parietal cortex samples from 31 sporadic late-onset AD cases (hAD), 16 autosomal dominant AD cases (APP or PSEN1 mutation carriers, hADAD) and 9 neuropathology-free controls (hCtrl). Cell-type-level differential expression results (hAD versus hCtrl and hADAD versus hCtrl) for microglia, astrocytes and oligodendrocytes were obtained from the authors on request as described in Brase et al.^33^. For each cell type, genes significantly dysregulated in hADAD (Benjamini–Hochberg-adjusted p < 0.05) were matched by gene symbol to the human orthologs of the genes differentially expressed between APP/PS1 and WT mice in the corresponding cell type, and genes with the same direction of change in both comparisons were retained (microglia, n = 14; astrocytes, n = 26; oligodendrocytes, n = 198, of which the 26 with the lowest hADAD adjusted p values were plotted). Human log fold changes reported by nebula (natural-log scale) were converted to log₂ fold changes for comparability with the mouse data. For these genes, log₂ fold changes for hAD versus hCtrl, hADAD versus hCtrl, APP/PS1 versus WT, AO versus WT and *Ogg1^-/-^* versus WT were plotted as heatmaps in R using ggplot2^83^, with genes ordered by APP/PS1 versus WT log₂FC. Benjamini–Hochberg-adjusted p values are shown for the human comparisons and nominal p values for the mouse comparisons.

### Cross-platform concordance analysis

Concordance of chromatin accessibility and transcriptional changes was assessed per genotype (*Ogg1^-/-^*, APP/PS1, AO) by comparing gene-level log₂ fold changes relative to WT from ATAC-seq, bRNA-seq and cell type-specific snRNA-seq (microglia, astrocytes, oligodendrocytes). For ATAC-seq, log₂ fold changes of promoter-annotated peaks were used and averaged when multiple peaks mapped to the same promoter. Log₂ fold changes were used irrespective of significance. The analysis was performed on the prevDEGs of each cell type. For each genotype and platform pair (ATAC-snRNA, ATAC-bRNA, bRNA-snRNA), direction agreement was quantified as Spearman’s rank correlation (ρ) of log₂ fold changes across genes with values in both platforms. Effect sizes were summarized per genotype and platform as the median absolute log₂ fold change across genes with values in all three platforms. Results were visualized as heatmaps in R using ggplot2^83^.

### Chord diagrams

Convergence of GO enrichment across datasets was visualized as chord diagrams using the circlize package^84^ in R. For each genotype (*Ogg1^-/-^*, APP/PS1, AO), top enriched GO biological process terms from the bRNA-seq, ATAC-seq and cell type-specific snRNA-seq (microglia, astrocytes, oligodendrocytes) analyses were compiled and grouped into three biological processes (immune response, synaptic signaling and myelination). Each dataset and each GO term is represented as a sector, and a chord connects a dataset to a GO term enriched in it, with chord width proportional to -log₁₀ FDR of the enrichment; terms not enriched in a dataset were assigned a value of 0 and therefore draw no chord. One diagram was generated per genotype.

### Statistical analysis

Statistical analyses were performed using GraphPad Prism (GraphPad Software, Inc., La Jolla, CA, USA; v. 10). Two-tailed Student’s *T*-tests were used for comparisons between two groups. For comparisons among multiple groups, ANOVA followed by an appropriate post-hoc test was used. Statistical significance was set at P < 0.05. Data are presented as mean ± s.e.m., unless stated otherwise. Data distributions were assumed to be normal but were not formally tested. All replicates represent independent biological replicates, with at least three biological replicates per experiment. Sample sizes were not determined using formal statistical methods. Exclusion criteria for behavioral experiments were predefined according to published guidance^85^.

## Data availability

All genomic sequencing data produced in the present project including bulk RNA-seq, ATAC-seq and snRNA-seq have been deposited in the NCBI GEO database under accession number GSE351382, GSE351381 and GSE351383, respectively.

## Acknowledgements

We thank the Norwegian Sequencing Centre in Oslo for the help to obtain Single nuclei RNA-seq data. Images were acquired at Cellular and Molecular Imaging Core Facility (CMIC), Norwegian University of Science and Technology (NTNU). CMIC is funded by the Faculty of Medicine and Health Sciences at NTNU, Central Norway Regional Health Authority and NALMIN II (NFR 322607). Mass spectrometry-based analyses were performed by the Proteomics and Modomics Experimental Core (PROMEC), Norwegian University of Science and Technology (NTNU) and The Central Norway Regional Health Authority. This facility is a member of the National Network of Advanced Proteomics Infrastructure (NAPI), which is funded by the Research Council of Norway INFRASTRUKTUR-program (NFR 295910).

## Contributions

AA, PS, MAE, KMA, CD, TO and MLS performed the experiments. AA, TS, PS, MAE, KMA, CD and MLS analyzed the data. NK provided access to human AD tissue samples. AA and KS designed the work. AA, TS, MB and KS interpreted the analysis. AA and KS wrote the manuscript. All authors participated in the review of the manuscript.

## Ethical approval

All experiments were approved by the Norwegian Animal Research Authority (NARA) and conducted in accordance with the laws and regulations controlling experimental procedures in live animals in Norway and the European Union’s Directive 86/609/EEC.

