## Supplementary material for "Ogg1 regulates context-dependent glial states in Alzheimer’s disease"

### Supplementary: Materials and Methods

#### Human post-mortem brain tissue processing

Post-mortem medial temporal lobe tissue was obtained from three healthy donors (HD; mean age, 77.3 ± 6.8 years) and three patients with Alzheimer's disease (hAD; mean age, 83.3 ± 11.6 years) from the UK brain bank. All donors were male. Approximately 30 mg of tissue was processed per sample. Each sample consisted of a mixture containing equal amounts of tissue from Brodmann area 28, Brodmann area 35 and the hippocampus. Snap-frozen tissue samples were homogenized in RLT buffer for 15 s at 3,000 rpm using a BeadBug™ 6 Six Microtube Homogenizer (Benchmark Scientific, Inc.).

#### Human gene expression analysis

Total RNA was isolated using the AllPrep DNA/RNA/Protein Mini Kit (Qiagen), according to the manufacturer's instructions. RNA quantity and purity was assessed using an ND-1000 spectrophotometer (NanoDrop Technologies). RNA was reverse transcribed using MultiScribe Reverse Transcriptase (Applied Biosystems) according to the manufacturer's protocol. Quantitative PCR (qPCR) was performed using Power SYBR Green PCR Master Mix (Applied Biosystems) on a StepOnePlus v2.3 Real-Time PCR System. Relative transcript levels were normalized to *ACTB* (β-actin), and gene expression was calculated using the ΔΔCt method. Primers used were as follows: *OGG1*, forward 5'-CTCCAACAACAACATCGCC-3', reverse 5'-GAGATGAGCCTCCACCTCTG-3'; *ACTB*, forward 5'-GTTACAGGAAGTCCCTTGCCATCC-3', reverse 5'-CACCTCCCCTGTGTGGACTTGGG-3'.

#### Targeted mass spectrometry

Protein was isolated using the AllPrep DNA/RNA/Protein Mini Kit (Qiagen), according to the manufacturer's instructions. Protein pellets were reconstituted in sodium deoxycholate (SDC) buffer consisting of 1% SDC, 10 mM tris (2-carboxyethyl) phosphine (TCEP), 40 mM chloroacetamide and 100 mM Tris-HCl pH 8.5, heated at 95 oC for 10 min, followed by sonication for 5 min. Proteins were digested overnight with trypsin (Thermo Scientific) at 37 oC, 850 rpm in a ThermoMixer (Thermo Scientific) and SDC was precipitated by formic acid. After centrifugation (1600xg), supernatant was desalted using home-made C18 stage tips and peptide concentration was measured using a NanodropOne C (A205 nm). Peptides were separated using an EASY-nLC 1200 UHPLC system (Thermo Scientific) coupled to a Q Exactive HF Orbitrap mass spectrometer (Thermo Scientific) as previously described[1]. PRM methods were designed and data were analyzed using Skyline (version 23.1)[2]. Relative OGG1 protein abundance was determined using the unique peptide LDLVLPSGQSFR.

**Supplementary figure 1**

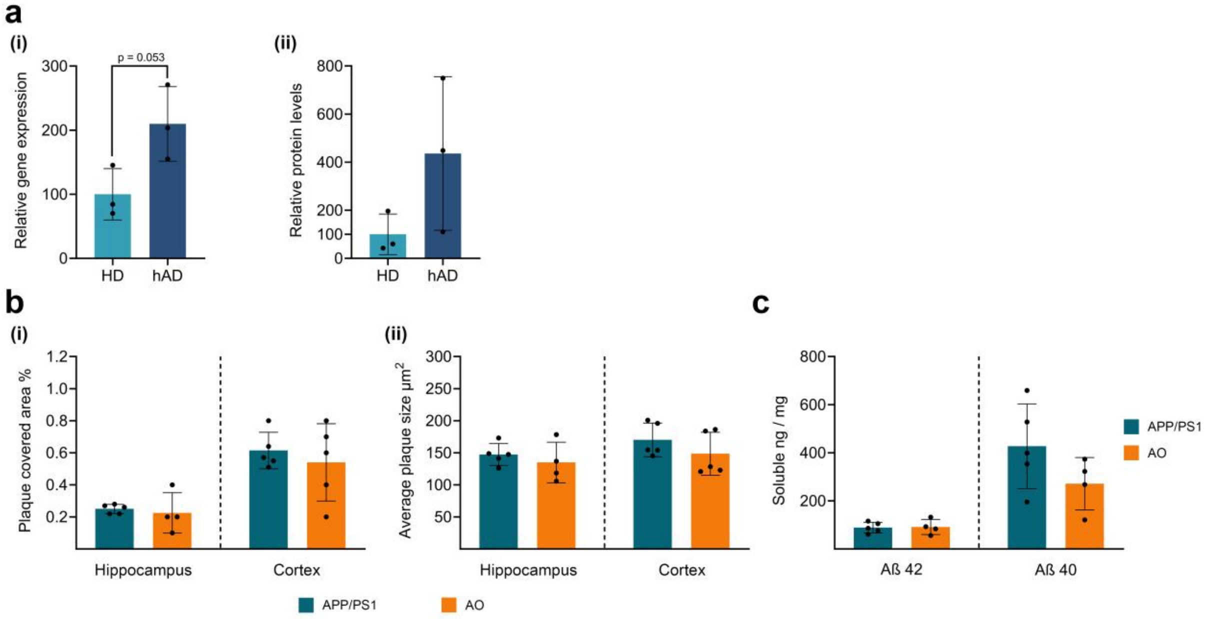


**Supplementary figure 1. Quantification of amyloid burden and oxidative DNA damage**

(a) (i) Relative *OGG1* mRNA expression measured by quantitative PCR (qPCR) and (ii) Relative OGG1 protein abundance measured by targeted mass spectrometry in post-mortem medial temporal lobe tissue from healthy donors (HD) and patients with Alzheimer's disease (hAD). Individual data points represent biological replicates; bars indicate mean ± SD. p values were determined using a two-tailed unpaired Student's *T*-test. (b) Quantification of (i) plaque-covered area and (ii) average plaque size in the hippocampus and cortex of APP/PS1 and AO mice (APP/PS1, n = 5; AO, n = 4). (c) Quantification of soluble Aβ42 and Aβ40 levels in whole-brain extracts measured by ELISA (APP/PS1, n = 5; AO, n = 4).

**Supplementary figure 2**

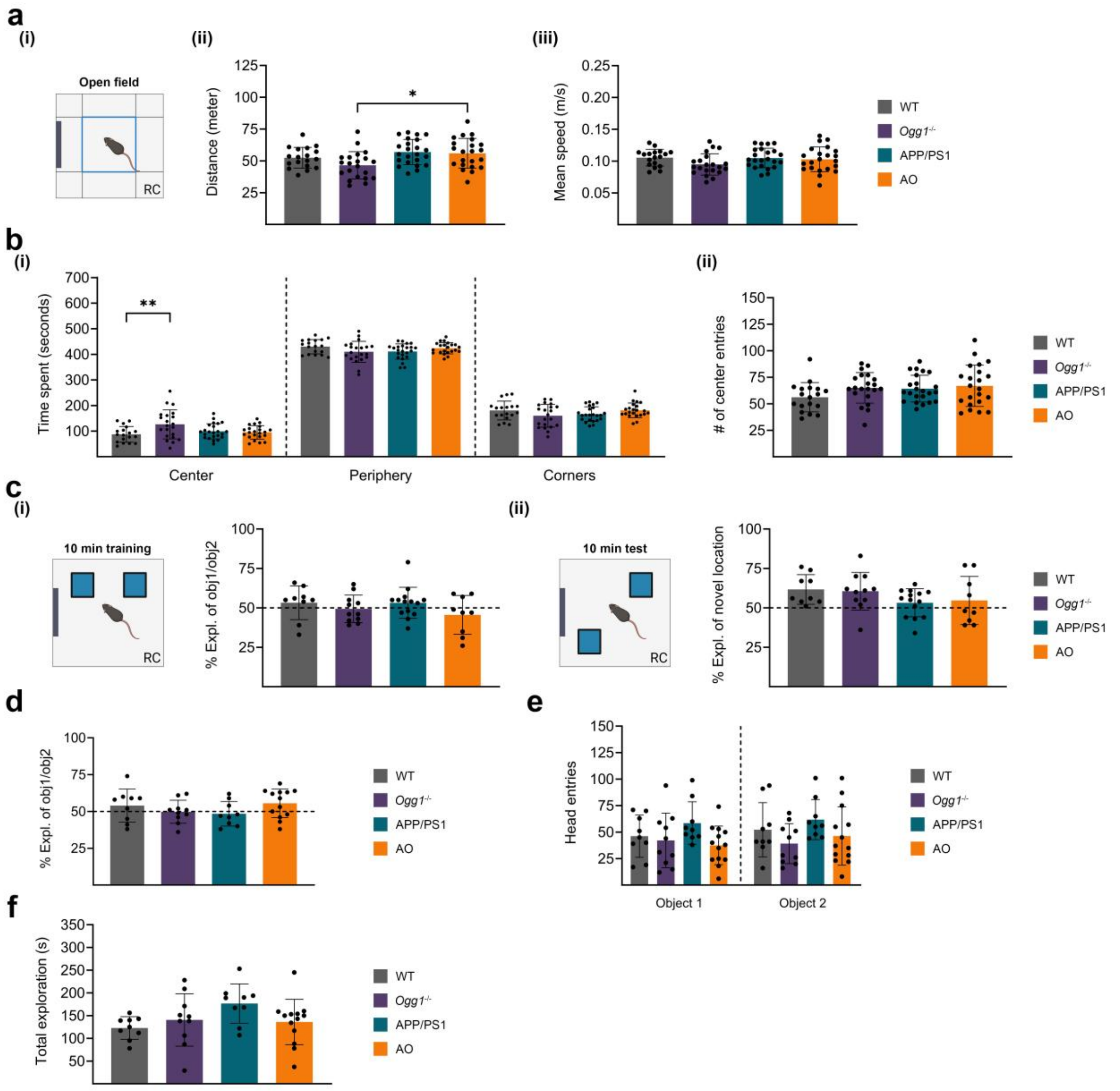


**Supplementary figure 2. Open field exploration and novel object location (NOL) behavior assessed by object preference and head entries**

(a) (i) Schematic of the open field test, indicating the releasing corner (RC), where mice were introduced into the arena. (ii) Total distance traveled and (iii) mean speed during the open field paradigm in WT, *Ogg1*$^{-/-}$, APP/PS1, and AO mice (WT, n = 18; *Ogg1*$^{-/-}$, n = 21; APP/PS1, n = 23; AO, n = 22). (b) Open field anxiety-related measures, including (i) time spent in the center versus peripheral zones and (ii) number of entries into the center area. (c) (i) Schematic of the novel object location (NOL) memory test, indicating the RC, where mice were introduced into the arena. (i) Preference for both object locations assessed during the training phase of the 1 h cohort in WT, *Ogg1*$^{-/-}$, APP/PS1, and AO mice (WT, n = 9; *Ogg1*$^{-/-}$, n = 11; APP/PS1, n = 14; AO, n = 9). (ii) Preference for the novel and familiar object locations assessed at the 1 h memory retention time point in the same genotypes. (d) Preference for both object locations assessed during the training phase of the 24 h cohort in WT, *Ogg1*$^{-/-}$, APP/PS1, and AO mice (WT, n = 9; *Ogg1*$^{-/-}$, n = 10; APP/PS1, n = 9; AO, n = 13). (e) Head entries toward each object assessed during the training phase of the 24 h NOL paradigm. (f) Total exploration time measured during the 24 h NOL test in WT, *Ogg1*$^{-/-}$, APP/PS1, and AO mice (WT, n = 9; Ogg1$^{-/-}$, n = 10; APP/PS1, n = 9; AO, n = 13). Data are presented as mean ± SEM, n values refer to the number of mice, *p < 0.05, **p < 0.01.

##### Supplementary figure 3

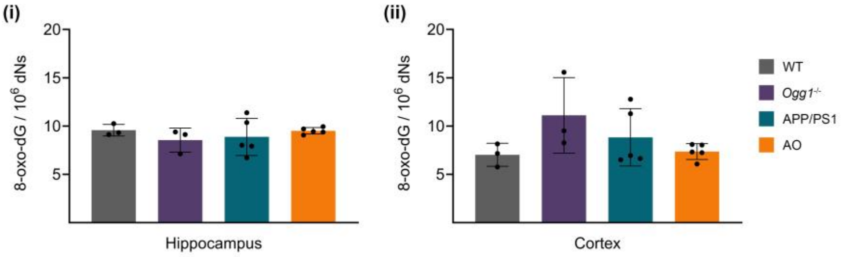


**Supplementary figure 3. Global oxidative DNA damage is unchanged across genotypes**

Quantification of oxidative DNA damage assessed by LC-MS/MS measurement of 8-oxo-7,8-dihydroguanine (8-oxoG) levels in (i) hippocampus and (ii) cortex of WT (n = 3), *Ogg1*$^{-/-}$ (n = 3), APP/PS1 (n = 5), and AO (n = 5) mice. Data are presented as mean ± SEM; n values refer to the number of mice.

##### Supplementary figure 4

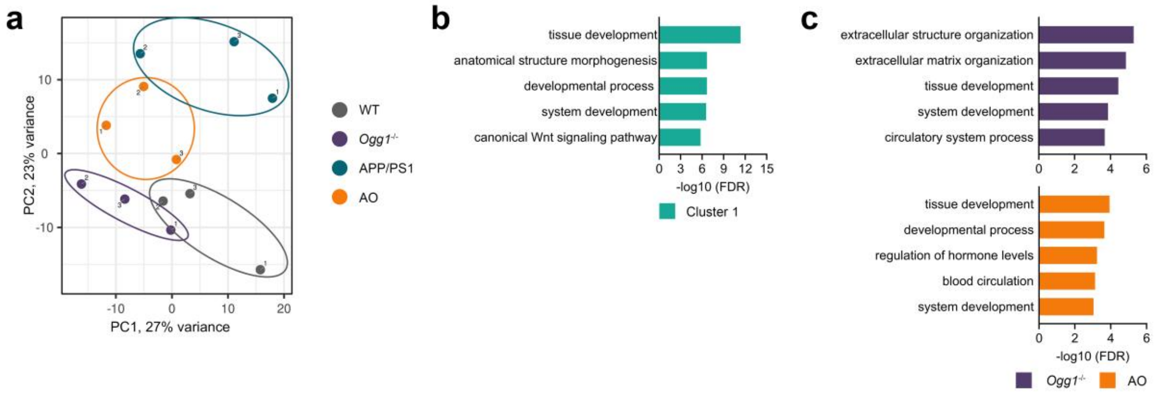


**Supplementary figure 4. Principal component analysis and functional annotation of bulk RNA-seq data**

(a) Principal component analysis (PCA) of total RNA-seq samples. (b) Gene ontology (GO) analysis of cluster 1 genes, with bar charts showing the top five biological process terms. (c) Enriched GO biological process terms for downregulated genes in *Ogg1*$^{-/-}$ and AO mice.

**Supplementary figure 5**

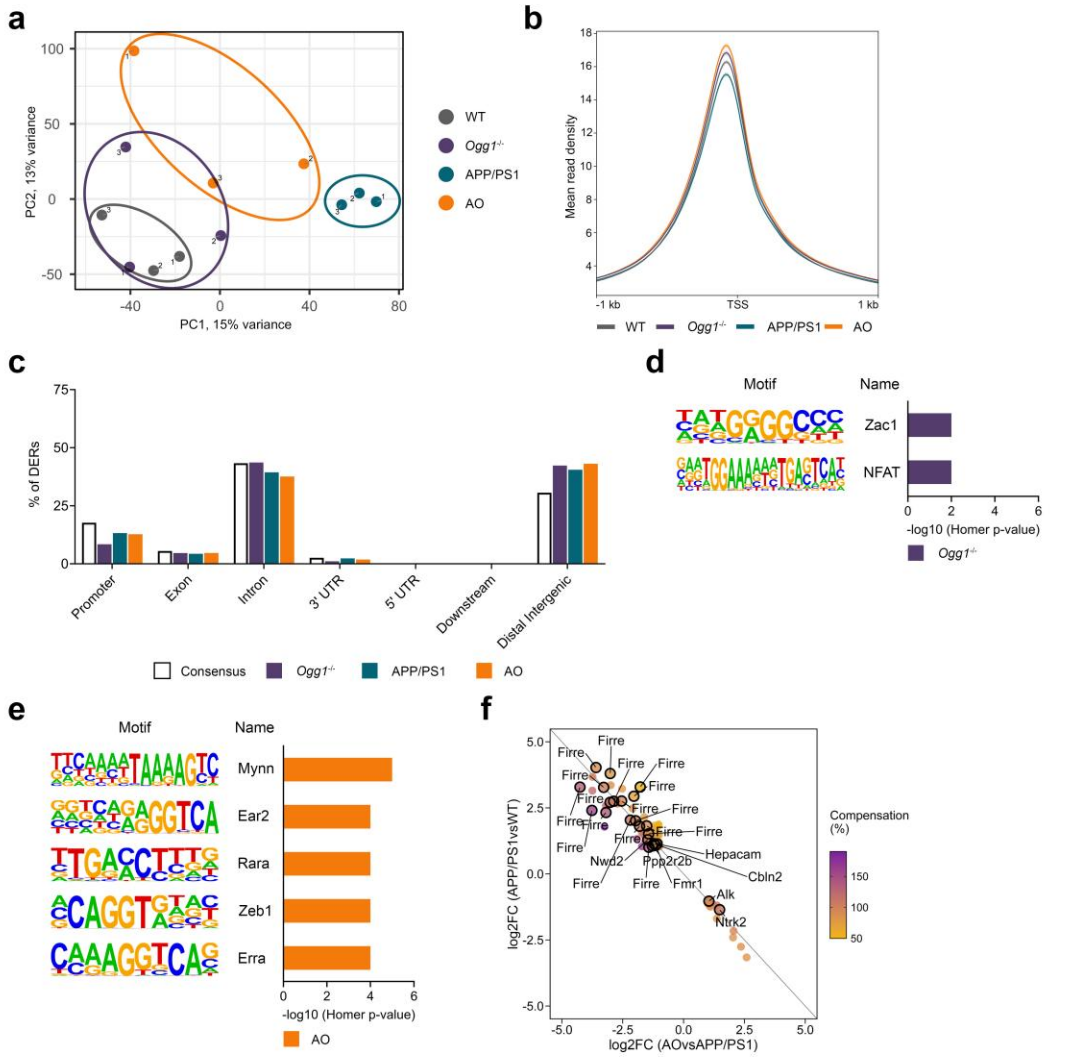


**Supplementary figure 5. Overview of ATAC-seq data, motif enrichment analysis, and genomic annotation**

(a) Principal component analysis (PCA) of ATAC-seq samples showing clustering of all biological replicates across genotypes. (b) Mean ATAC-seq signal intensity profile centered on transcription start sites (TSS). (c) Genomic annotation of all differentially accessible regions (DARs) identified by ATAC-seq. Consensus peaks were defined as regions detected in at least two of the three biological replicates. (d-e) DNA motif enrichment analysis of (d) *Ogg1*$^{-/-}$-specific and (e) AO-specific DARs, combining gained and lost regions. (f) Scatter plot showing the relationship between chromatin accessibility changes in APP/PS1 versus WT and AO versus APP/PS1 mice. Each dot represents a preventative differentially accessible region (prevDAR). The color gradient indicates the compensation rate (%), calculated as the ratio of the $\log_2$ fold changes in AO versus APP/PS1 to that in APP/PS1 versus WT, with higher values indicating greater prevention of AD-associated changes in chromatin accessibility. Regions outlined in black indicate selected prevDARs.

**Supplementary figure 6**

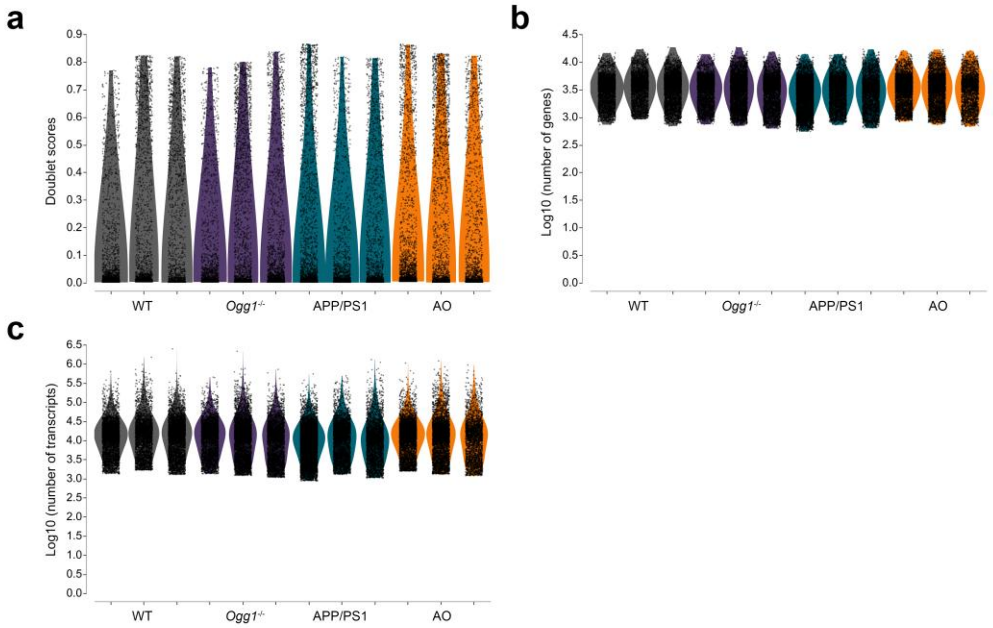


**Supplementary figure 6. Quality control metrics for single-nuclei RNA sequencing data**

(a-c) Violin plot displaying (a) doublet scores for individual nuclei, (b) number of detected genes per nucleus and (c) number of detected transcripts per nucleus across all three biological replicates per genotype.

**Supplementary figure 7**

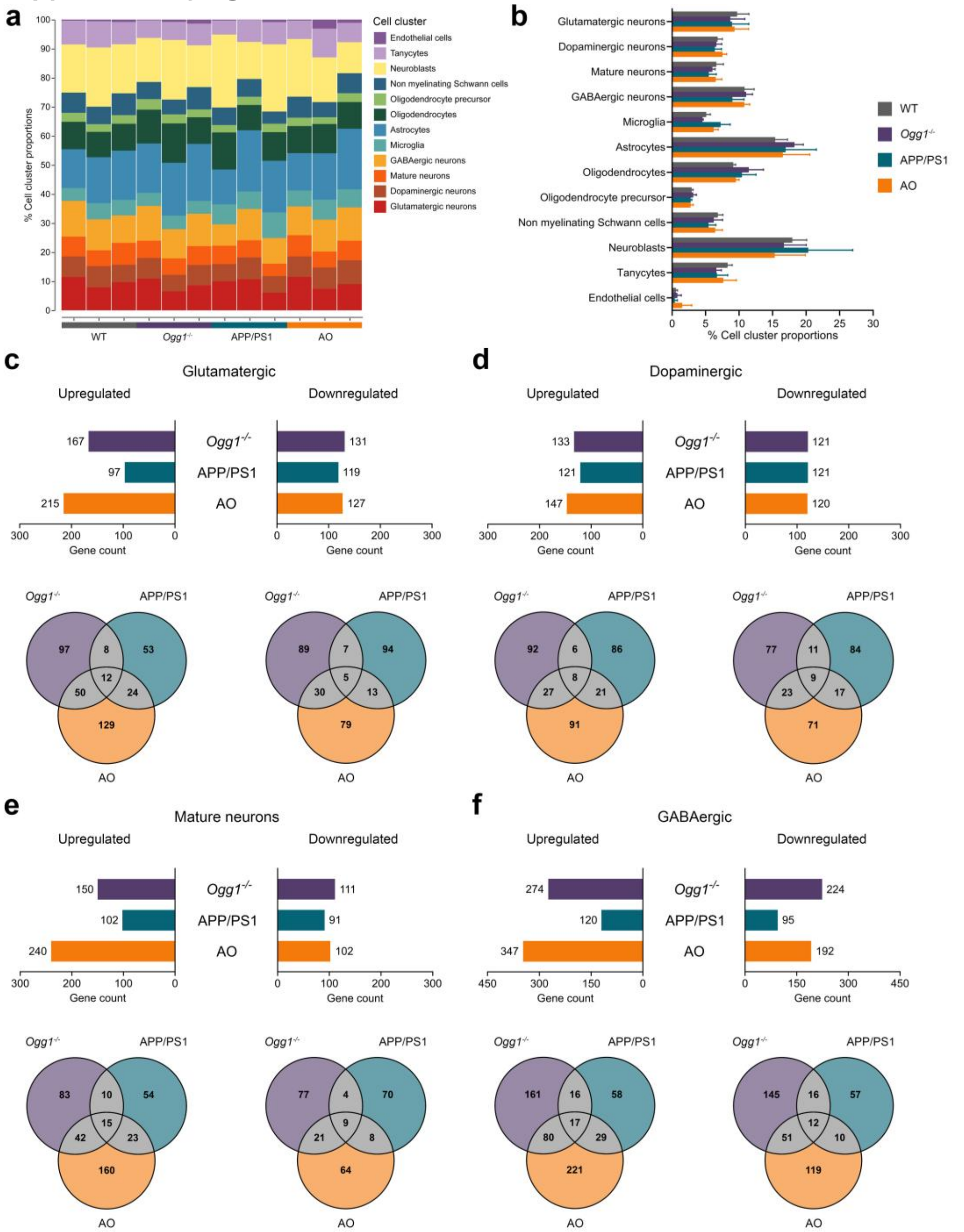


**Supplementary figure 7. Cell cluster distribution and neuron subtype-specific differential expression in snRNA-seq data**

(a) Proportional distribution of all cell clusters shown for each of the three individual biological replicates per genotype. (b) Proportional distribution of all cell clusters summarized as a bar graph across genotypes. Data are presented as mean ± SEM. (c–f) Neuron subtype-specific differential gene expression analysis from snRNA-seq data. Bar graphs illustrate the number of differentially expressed genes in glutamatergic, dopaminergic, mature, and GABAergic neurons, respectively for *Ogg1*$^{-/-}$, APP/PS1, and AO compared to WT ($|\log_2$ fold change$| \geq 0.5$, $p < 0.05$). Venn diagrams depict unique and shared upregulated and downregulated genes across genotypes for each neuronal subtype.

**Supplementary figure 8**

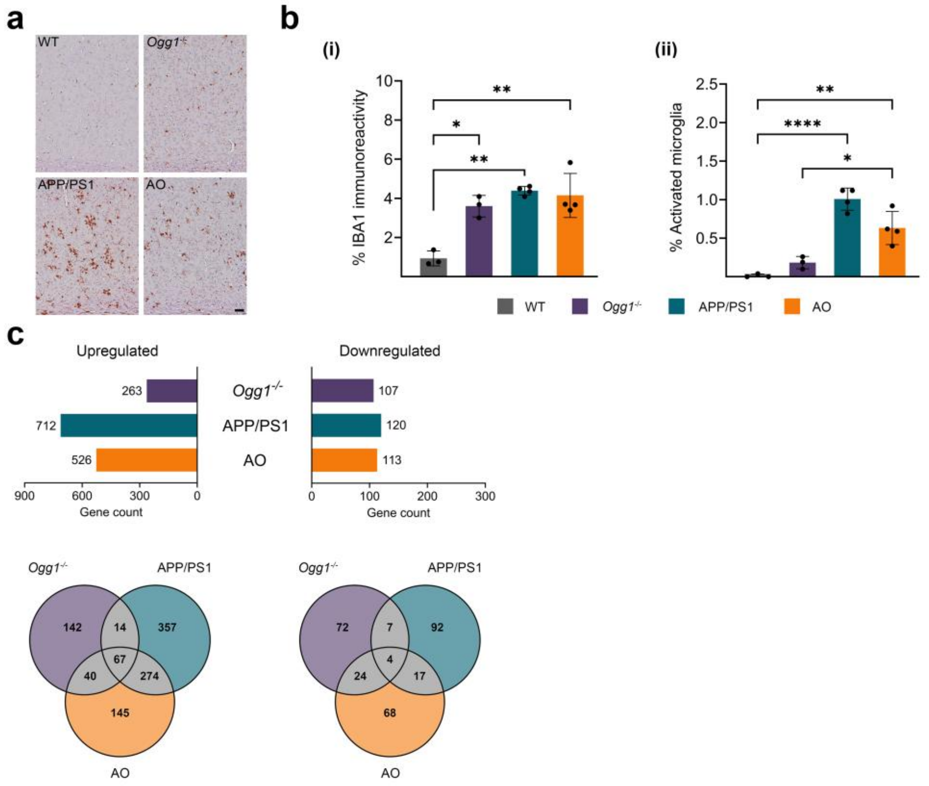


**Supplementary figure 8. Microglia characterization by immunostaining and snRNA-seq**

(a) Representative images of IBA1 immunostaining in the cortex of 6 months old mice across all genotypes. Scale bar: 20 μm.(b) (i) Quantification of cortical IBA1 immunoreactivity and (ii) Relative proportion of activated cortical IBA1 positive microglia of WT, *Ogg1*$^{-/-}$, APP/PS1, and AO mice at 6 months of age. Data are presented as mean ± SEM, n values refer to the number of mice, *$p < 0.05$, **$p < 0.01$, ***$p < 0.001$, ****$p < 0.0001$. (c) Microglia-specific differential gene expression analysis from snRNA-seq data. Bar graphs illustrate the number of differentially expressed genes for *Ogg1*$^{-/-}$, APP/PS1, and AO compared to WT ($|\log_2$ fold change$| \geq 0.5$, $p < 0.05$). Venn diagrams depict unique and shared upregulated and downregulated genes across genotypes.

**Supplementary figure 9**

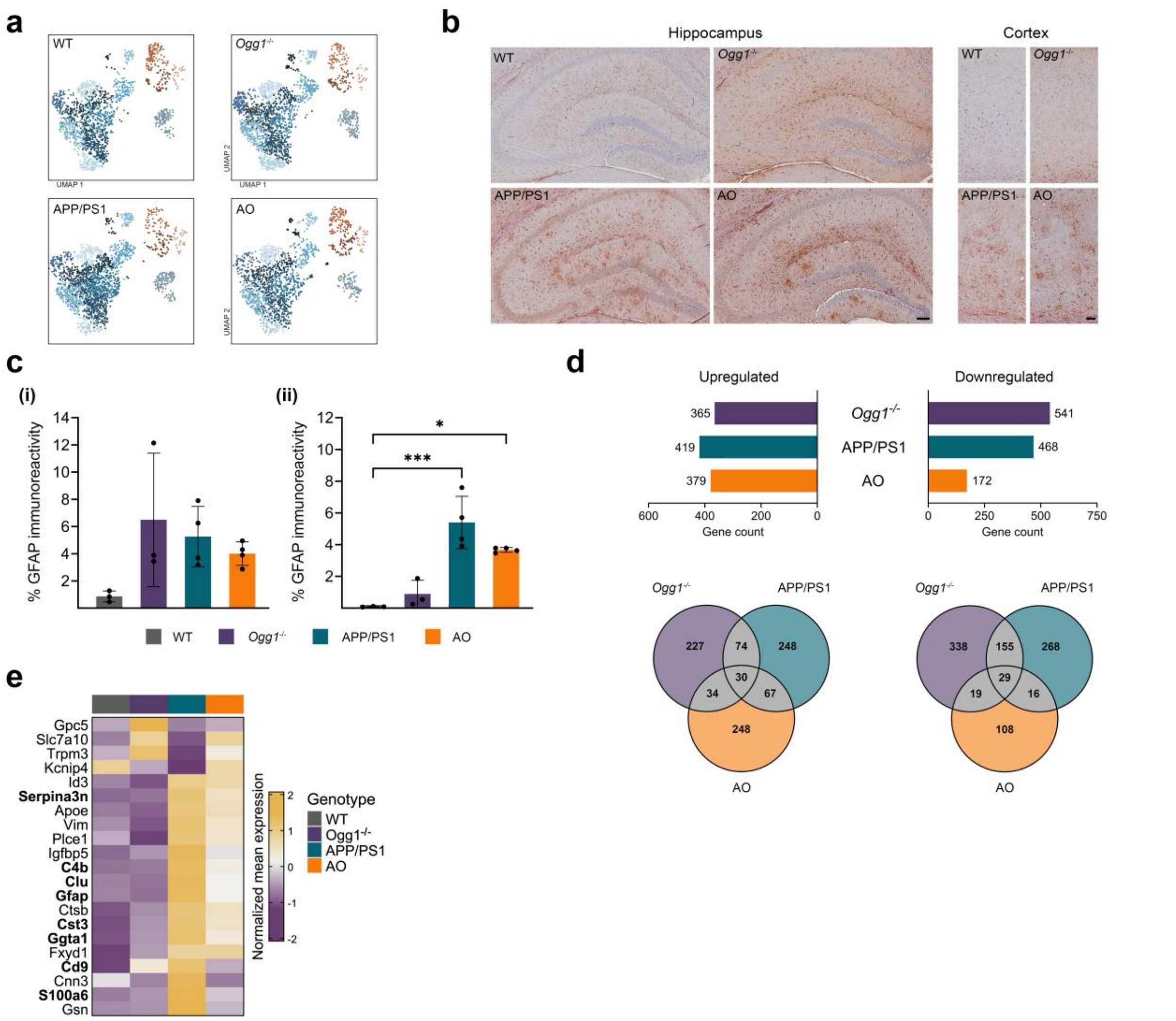


**Supplementary figure 9. Immunohistochemical and transcriptomic snRNA-seq analysis of astrocytes**

(a) UMAP representations of astrocyte nuclei from snRNA-seq shown for each genotype, revealing transcriptomically defined subclusters annotated by gene ontology (GO) biological process enrichment. (b) Representative images of GFAP immunostaining in the hippocampus and cortex of 6 months old mice across all genotypes. Scale bars: 100 µm (hippocampus) and 20 µm (cortex). (c) Quantification of GFAP immunoreactivity in the (i) hippocampus and (ii) cortex of WT, Ogg1$^{-/-}$, APP/PS1, and AO mice at 6 months of age. Data are presented as mean ± SEM, n values refer to the number of mice, *$p < 0.05$, **$p < 0.01$, ***$p < 0.001$. (d) Astrocyte-specific differential gene expression analysis from single-nucleus RNA sequencing (snRNA-seq) data. Bar graphs illustrate the number of differentially expressed genes for Ogg1$^{-/-}$, APP/PS1, and AO compared to WT ($|\log_2$ fold change$| \geq 0.5$, $p < 0.05$), with Venn diagrams depicting unique and shared upregulated and downregulated genes across genotypes. (e) Heatmap showing the mean expression of disease-associated astrocyte genes, as previously reported[3], across genotypes in this dataset.

**Supplementary figure 10**

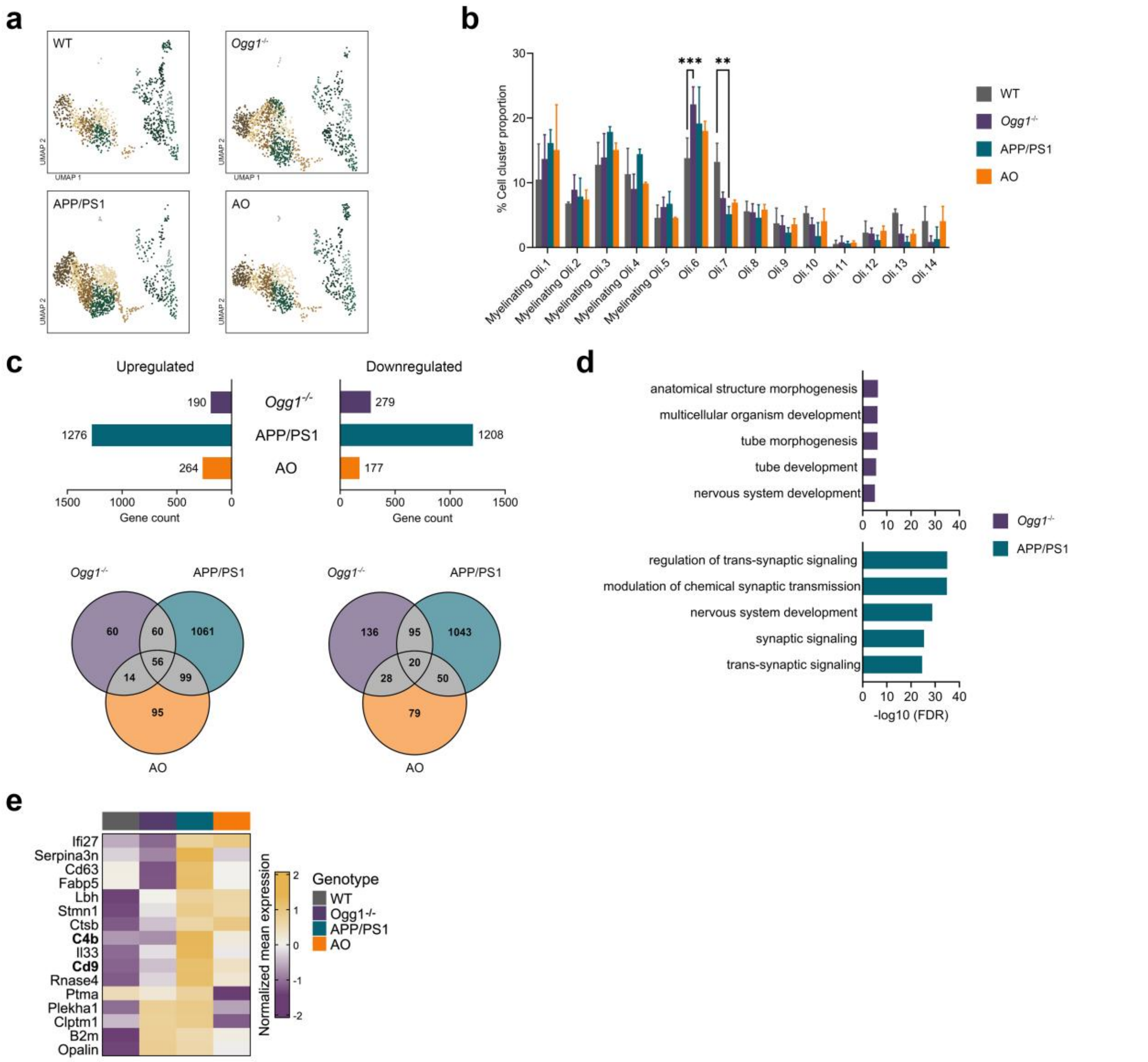


**Supplementary figure 10. Oligodendrocyte subclustering and differential gene expression analysis by snRNA-seq** (a) UMAP representations of oligodendrocyte nuclei from snRNA-seq shown for each genotype, revealing transcriptomically defined subclusters annotated by gene ontology (GO) biological process enrichment. (b) Percentage distribution of nuclei among the oligodendrocyte subclusters. Data are presented as mean ± SEM, n values refer to the number of mice, *p < 0.05, **p < 0.01, ***p < 0.001. (c) Oligodendrocyte-specific differential gene expression analysis of snRNA-seq data. Bar graphs illustrate the number of differentially expressed genes for *Ogg1*$^{-/-}$, APP/PS1, and AO compared to WT ($|\log_2$ fold change$| \geq 0.5$, $p < 0.05$), with Venn diagrams depicting unique and shared upregulated and downregulated genes across genotypes. (d) Enriched GO biological process terms for unique downregulated genes in *Ogg1*$^{-/-}$ and APP/PS1. (e) Heatmap showing the mean expression of disease-associated oligodendrocyte genes, as previously reported[4], across genotypes in this dataset.
